# Accounting for motion in resting-state fMRI: What part of the spectrum are we characterizing in autism spectrum disorder?

**DOI:** 10.1101/2022.01.12.476077

**Authors:** Mary Beth Nebel, Daniel E. Lidstone, Liwei Wang, David Benkeser, Stewart H. Mostofsky, Benjamin B. Risk

**Author notes:** 716 N Broadway, Baltimore, MD 21205 *Email address:* (Mary Beth Nebel).

## Abstract

The exclusion of high-motion participants can reduce the impact of motion in functional Magnetic Resonance Imaging (fMRI) data. However, the exclusion of high-motion participants may change the distribution of clinically relevant variables in the study sample, and the resulting sample may not be representative of the population. Our goals are two-fold: 1) to document the biases introduced by common motion exclusion practices in functional connectivity research and 2) to introduce a framework to address these biases by treating excluded scans as a missing data problem. We use a study of autism spectrum disorder in children without an intellectual disability to illustrate the problem and the potential solution. We aggregated data from 545 children (8-13 years old) who participated in resting-state fMRI studies at Kennedy Krieger Institute (173 autistic and 372 typically developing) between 2007 and 2020. We found that autistic children were more likely to be excluded than typically developing children, with 28.5% and 16.1% of autistic and typically developing children excluded, respectively, using a lenient criterion and 81.0% and 60.1% with a stricter criterion. The resulting sample of autistic children with usable data tended to be older, have milder social deficits, better motor control, and higher intellectual ability than the original sample. These measures were also related to functional connectivity strength among children with usable data. This suggests that the generalizability of previous studies reporting naïve analyses (i.e., based only on participants with usable data) may be limited by the selection of older children with less severe clinical profiles because these children are better able to remain still during an rs-fMRI scan. We adapt doubly robust targeted minimum loss based estimation with an ensemble of machine learning algorithms to address these data losses and the resulting biases. The proposed approach selects more edges that differ in functional connectivity between autistic and typically developing children than the naïve approach, supporting this as a promising solution to improve the study of heterogeneous populations in which motion is common.

## 1. Introduction

Resting-state functional magnetic resonance imaging (rs-fMRI) relies on spontaneous, interregional correlations in blood-oxygen-level-dependent signal fluctuations, termed functional connectivity, to characterize brain organization (Biswal et al., 1995). A fundamental challenge in rs-fMRI-based research is to separate the signal reflecting neural activity from a combination of unstructured thermal noise and spatiotemporally structured signals of non-interest. Participant head motion is problematic because even sub-millimeter movements can introduce spatially variable artifacts that are challenging to correct during postprocessing (Power et al., 2012; van Dijk et al., 2012; Satterthwaite et al., 2012). Post-acquisition motion quality control (QC) procedures involve two stages: 1) elimination of scans with gross motion (scan exclusion); and 2) minimization of artifacts due to subtle motion (de-noising). Guidelines for removing motion-corrupted rs-fMRI data have been proposed (Satterthwaite et al., 2013; Parkes et al., 2018; Power, 2017), and many post-acquisition cleaning procedures have been developed (Satterthwaite et al., 2013; Power et al., 2014; Muschelli et al., 2014; Pruim et al., 2015; Mejia et al., 2017; Power et al., 2020). However, this work has focused on maximizing rs-fMRI data quality. The impact of scan exclusion on the study sample composition and selection bias has been largely unexamined.

Motion is particularly common in pediatric and clinical populations (Fassbender et al., 2017; Greene et al., 2018). The focus on maximizing rs-fMRI data quality has been driven by a concern that if motion artifacts are not rigorously cleaned from the data, they may introduce spurious functional connectivity differences between groups of interest. For example, autism spectrum disorder (ASD) is a neurodevelopmental condition affecting approximately 1 in 44 children in the United States that is characterized by impairments in social and communicative abilities as well as restricted interests and repetitive behaviors (Maenner et al., 2021; American Psychiatric Association, 2013). The ‘connectivity hypothesis’ of autism claims that short-range connections are increased at the expense of long-range connections within the brain (for a review, see Vasa et al. (2016)). However, sub-millimeter motion-related artifacts often mimic this pattern. High-motion participants show stronger correlations between nearby brain locations and weaker correlations between distant brain regions compared to low-motion participants, even after controlling for motion in multiple modeling steps (Power et al., 2012; van Dijk et al., 2012; Satterthwaite et al., 2012). ASD functional connectivity studies have found conflicting patterns of widespread hypoconnectivity, hyperconnectivity, and mixtures of the two (Di Martino et al., 2011; Supekar et al., 2013; Rudie et al., 2013; Keown et al., 2013; Dajani and Uddin, 2016; Lombardo et al., 2019). Moreover, studies using stricter motion QC have reported largely typical patterns of functional connectivity (Tyszka et al., 2014), suggesting that motion artifacts may have contributed to discrepancies in the literature (Deen and Pelphrey, 2012).

Exclusion of high-motion participants may help alleviate motion artifacts in functional connectivity estimates but may also introduce a new problem by systematically altering the study population. Implementation of scan exclusion guidelines can lead to drastic reductions in sample size. For instance, in a study examining the impact of motion artifact de-noising procedures on predictions of brain maturity from rs-fMRI data, Nielsen et al. (2019) excluded 365 of 487 participants between 6 and 35 years of age due to excessive head motion. Applying similarly stringent scan exclusion criteria to rs-fMRI data from the Adolescent Brain Cognitive Development (ABCD) study, Marek et al. (2019) excluded 40% of participants despite efforts by the ABCD study to track head motion in real-time (Dosenbach et al., 2017) to ensure a sufficient amount of motion-free data would be collected from each participant (Casey et al., 2018). One strategy for balancing the need to rigorously clean the data with the cost of excluding participants has been to use less stringent scan exclusion criteria and then examine the effect of diagnosis in a linear model controlling for age and summary measures of between-frame head motion (e.g., Di Martino et al. 2014). However, the possibility of introducing selection bias following scan exclusion remains.

In studies in which the outcome is missing for some participants, the difference in the mean between two groups calculated from observed outcomes may be biased if data are not randomly missing (Hernan and Robins, 2020). In rs-fMRI studies, naïve estimators of group-level functional connectivity based only on participants with usable rs-fMRI data may be biased if scan exclusion changes the distribution of participant characteristics related to functional connectivity. In the case of ASD, studies excluding high-motion participants have reported functional connectivity differences between autistic and typically developing children, as well as associations between functional connectivity strength and the severity of motor and social skill deficits (Uddin et al., 2013; Lake et al., 2019; Wymbs et al., 2021; D’Souza et al., 2021), but these studies did not examine the impact of scan exclusion on the composition of the study sample with usable data. The graph in Figure 1 illustrates how excluding high-motion participants (Δ = 0) could obscure the relationship between a diagnosis of ASD (*A*) and functional connectivity (*Y*) by changing the joint distribution of diagnosis and a covariate related to symptom severity (*W*). Bias can arise from a lack of exchangeability, as children with usable data differ from those with unusable data. The term “‘confounding bias” is sometimes used to describe bias that arises when a predictor and outcome share a common cause (e.g., Δ → *W* and Δ → *Y*). The term “selection bias” or “Berksonian bias” may be used to describe bias from conditioning on common effects (e.g., *W* → Δ and *Y* → Δ). These concepts often overlap (Lash et al., 2021). In both cases, the key source of bias is a violation of exchangeability (Hernan and Robins, 2020). In Fig. 1, we describe the bias as selection bias, since it can be viewed as originating from a study that selects children that pass motion QC. If autistic children with usable rs-fMRI data are phenotypically more similar to typically developing children than those that were excluded, observed group differences may be reduced relative to group differences if we were able to collect usable rs-fMRI data from all participants.

**Figure 1:**
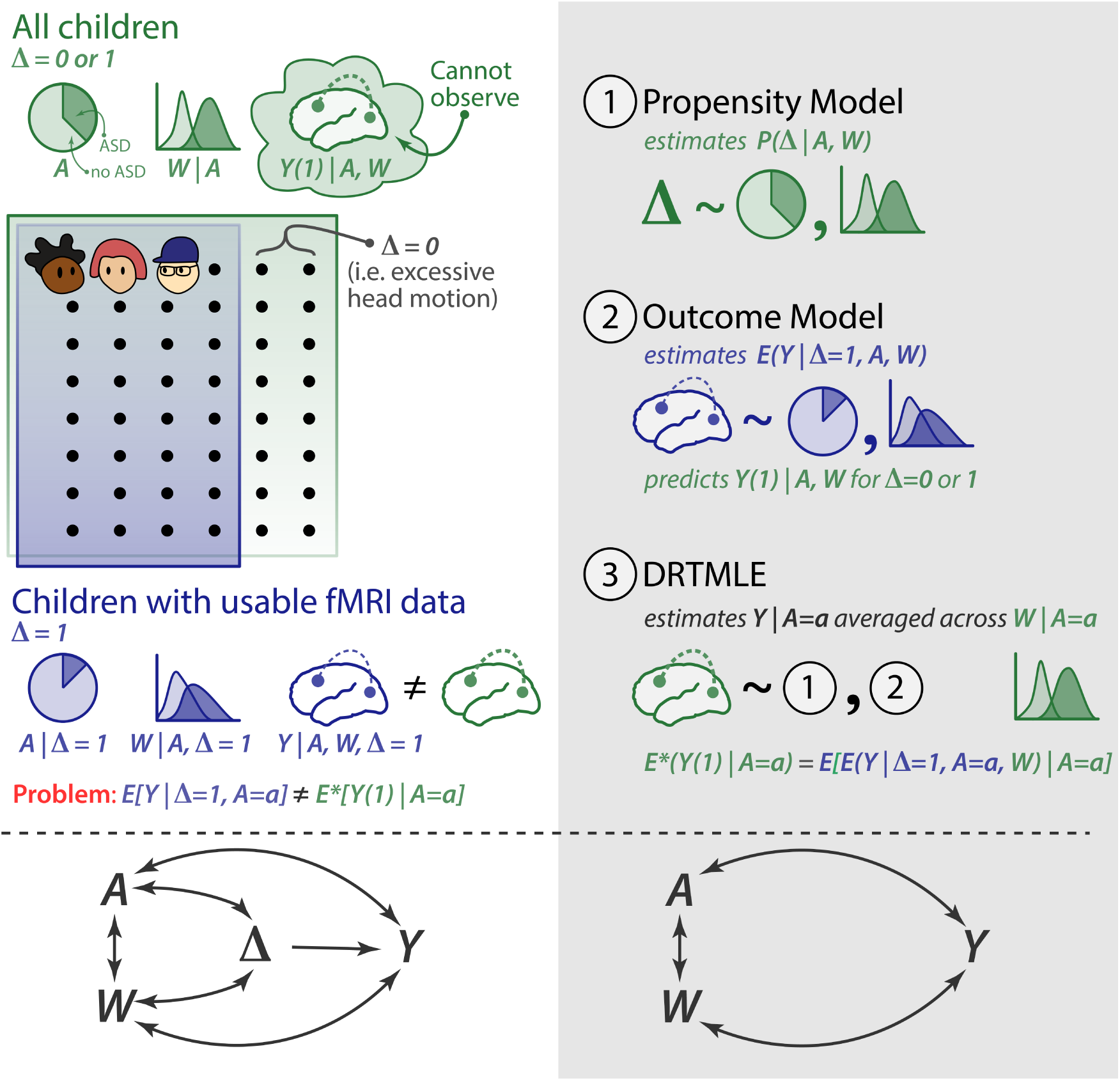
Scan exclusion may induce selection bias. *A* indicates diagnosis, where *A* = 0 (lighter shading) represents the typically developing group and *A* = 1 (darker shading) represents the autism spectrum disorder (ASD) group. *W* represents a covariate that reflects symptom severity; Δ indicates resting-state fMRI usability, where Δ=1 is usable and Δ=0 is unusable. *Y* is the functional connectivity between two brain regions, and *Y* (1) is the possibly counter fact functional connectivity from a usable fMRI scan. *E*^*^() denotes an expectation with respect to the probability measure of *Y* (1)*, A, W* . Additional details are in Section 2.3.1. (Left panel) Children with usable resting-state fMRI data (in purple) may systematically differ from all enrolled children (in green). The distribution of symptom severity *W* differs between children with usable and unusable fMRI data (*W* Δ). *W* is related to functional connectivity (*W Y*). Additionally, there are associational effects between ASD and functional connectivity (*A Y*) and between ASD and the symptom severity covariate (*A W*). Under these conditions, naïve estimators of group-level functional connectivity based only on participants with usable data may be biased. (Right panel) We propose to address this bias using doubly robust targeted minimum loss based estimation (DRTMLE), which involves three steps. 1. Fit the propensity model. 2. Fit the outcome model, which predicts functional connectivity from the covariates for participants with usable rs-fMRI data. Then use this model to predict functional connectivity for both usable and unusable participants. 3. Apply the DRTMLE algorithm, which uses the inverse probability of usability from step 1 and predictions of functional connectivity for all subjects (usable and unusable) from step 2 to break the pathways *A* ↔ Δ, *W* ↔ Δ, and Δ → *Y* .

In this study, we first describe our motivating dataset, an aggregation of phenotypic and rs-fMRI data from 173 autistic children without an intellectual disability and 372 typically developing children who participated in one of several neuroimaging studies at Kennedy Krieger Institute (KKI) between 2007 and 2020. We then explore the impact of commonly used head motion exclusion criteria on the composition of the sample of participants with usable rs-fMRI data, so that we can better understand what part of the spectrum we are characterizing after accounting for motion. Next we introduce a method for estimating functional connectivity adjusting for the observed sampling bias following participant exclusion due to motion QC, which we call the deconfounded group difference (Figure 1, grey panel). We propose to treat the excluded rs-fMRI scans as a missing data problem. We use an ensemble of machine learning algorithms to estimate the relationship between behavioral phenotypes and rs-fMRI data usability, which is called the propensity model, and between behavioral phenotypes and functional connectivity, which is called the outcome model. The propensity and outcome models are then used in the doubly robust targeted minimum loss based estimation (DRTMLE) of the deconfounded group difference (Benkeser et al., 2017; van der Laan and Rose, 2011; van der Laan et al., 2007). We apply this approach to estimate the deconfounded group difference between autistic children without intellectual disabilities and typically developing children in the KKI dataset and compare our findings to the naïve approach. Finally, we discuss the costs and benefits of motion quality control and our proposed solution.

## 2. Methods

### 2.1. Dataset

#### 2.1.1. Study Population

Our initial cohort is an aggregate of 545 children between 8- and 13-years old who participated in one of several neuroimaging studies at Kennedy Krieger Institute (KKI) between 2007 and 2020. Participants included 173 autistic children without an intellectual disability (148 boys) and 372 typically developing children (258 boys); rs-fMRI scans and a limited set of phenotypic data from 266 of these children (78 with ASD) were previously shared with the Autism Brain Imaging Data Exchange (ABIDE) (Di Martino et al., 2014, 2017). Participants were recruited through local schools, community-wide advertisement, volunteer organizations, medical institutions, and word of mouth. The data collecting studies were all approved by the Johns Hopkins University School of Medicine Institutional Review Board. After providing a complete study description, informed consent was obtained from a parent/guardian prior to the initial phone screening; written informed consent and assent were obtained from the parent/guardian and the child, respectively, upon arrival at the initial laboratory visit.

The following exclusion criteria were defined by the data-collecting studies. Children were ineligible to participate if their full scale intelligence quotient (FSIQ) from the Wechsler Intelligence Scale for Children, Fourth or Fifth Edition (WISC-IV or WISC-V; (Wechsler, 2003)) was less than 80 and they scored below 65 on 1) the Verbal Comprehension Index and 2) the Perceptual Reasoning Index (WISC-IV) or the Visual Spatial Index and the Fluid Reasoning Index (WISC-V), depending on which version of the WISC was administered. Children were also excluded if they had a) a history of a definitive neurological disorder, including seizures (except for uncomplicated brief febrile seizures), tumor, lesion, severe head injury, or stroke, based on parent responses during an initial phone screening; b) a major visual impairment; or c) conditions that contraindicate or make it challenging to obtain MRI data (e.g., cardiac pacemaker, surgical clips in the brain or blood vessels, or dental braces).

A diagnosis of ASD was determined using the Autism Diagnostic Observation Schedule-Generic (Lord et al., 2000) or the Autism Diagnostic Observation Schedule, Second Edition (ADOS-2) (Lord and Jones, 2012), depending on the date of enrollment. Diagnosis was verified by a board-certified child neurologist (SHM) with more than 30 years of experience in the clinical assessment of autistic children. Autistic children were excluded if they had identifiable causes of autism (e.g., fragile X syndrome, Tuberous Sclerosis, phenylketonuria, congenital rubella), documented history of prenatal/perinatal insult, or showed evidence of meeting criteria for major depression, bipolar disorder, conduct disorder, or adjustment disorder based on parent responses during an initial phone screening. Within the ASD group, a secondary diagnosis of attention deficit hyperactivity disorder (ADHD) was determined using the DSM-IV or DSM–5 (APA, 2000; APA, 2013) criteria and confirmed using a structured parent interview, either the Diagnostic Interview for Children and Adolescents-IV (DICA-IV; Reich (2000)) or the Kiddie Schedule for Affective Disorders and Schizophrenia for School-Age Children (K-SADS; Kaufman et al. (2013)), as well as parent and teachers versions of the Conners-Revised (Conners, 1999) or the Conners-3 Rating Scale (Conners, 2008), and parent and teacher versions of the DuPaul ADHD Rating Scale (DuPaul et al., 1998). To be classified as having comorbid ASD and ADHD (ASD+ADHD), a child with ASD had to receive one of the following: 1) a t-score of 60 or higher on the inattentive or hyperactive subscales of the Conners’ Parent or Teacher Rating Scale, or 2) a score of 2 or 3 on at least 6 of 9 items on the Inattentive or Hyperactivity/Impulsivity scales of the ADHD Rating Scale-IV (DuPaul et al., 1998). Diagnosis was verified by a board-certified child neurologist (SHM) or clinical psychologist with extensive experience in the clinical assessment of children with ADHD. Children taking stimulant medications were asked to withhold their medications the day prior to and the day of their study visit to avoid the effects of stimulants on cognitive, behavioral, and motor measures.

Children were excluded from the typically developing group if they had a first-degree relative with ASD, if parent responses to either the DICA-IV or for more recent participants, the K-SADS, revealed a history of a developmental or psychiatric disorder, except for simple phobias, or if they scored above clinical cut-offs on the parent and teacher versions of the Conners’ and ADHD Rating Scales.

The Hollingshead Four-Factor Index was used to generate a composite score of family socioeconomic status (SES) for each participant based on each parent’s education, occupation, and marital status (Hollingshead, 1975). Higher scores reflect higher SES.

#### 2.1.2. Phenotypic Assessment

The severity of core ASD symptoms was quantified within the ASD group using scores from the ADOS or the ADOS-2 calibrated to be comparable across instrument versions (Hus et al., 2014). We focus on the ADOS/ADOS-2 Comparable Total Score, hereafter refered to as ADOS; higher total scores indicate more severe ASD symptoms. These semi-structured ASD observation schedules are rarely administered to control participants; they were not designed to characterize meaningful variability in unaffected individuals, and scores are usually equal or close to zero in typically developing children. However, ASD-like traits vary among non-clinical individuals, with those meeting criteria for a diagnosis of ASD falling at one extreme of a spectrum encompassing the population at large. To supplement ADOS information, parent and teacher responses to the Social Responsiveness Scale (SRS) questionnaire (Constantino and Todd, 2003) or the SRS-2 (Constantino and Gruber, 2012) were also used. The SRS asks a respondent to rate a child’s motivation to engage in social interactions and their ability to recognize, interpret, and respond appropriately to emotional and interpersonal cues. The SRS yields a total score ranging between 0 and 195, with a higher total score indicating more severe social deficits. Total raw scores were averaged across respondents.

We also quantified the severity of ADHD symptoms using parent responses to the DuPaul ADHD Rating Scale (DuPaul et al., 1998) due to the high comorbidity of ASD and ADHD (Simonoff et al., 2008) and previous reports associating in-scanner movement with ADHD-like traits (Kong et al., 2014). The DuPaul ADHD Rating Scale asks a caregiver to rate the severity of inattention and hyperactivity/impulsivity symptoms over the last six months and yields a total raw score as well as two domain scores: inattention and hyper-activity/impulsivity. Our analyses focus on the two domain scores; higher DuPaul scores indicate more severe symptoms.

In addition to ASD and ADHD trait severity, basic motor control was examined using the Physical and Neurological Exam for Subtle Signs (PANESS), as the children were, in effect, asked to complete a motor task by remaining as still as possible during the scan. The PANESS assesses basic motor control through a detailed examination of subtle motor deficits, including overflow movements, involuntary movements, and dysrhythmia (Denckla, 1985), which also allows for the observation of handedness. We focused on total motor overflow as our primary measure of motor control derived from the PANESS. Motor overflow is a developmental phenomenon defined as unintentional movements that mimic the execution of intentional movements. Motor overflow is common in early childhood and typically decreases as children age into adolescence. Excessive degree and abnormal persistence of motor overflow is thought to reflect an impaired capacity to inhibit unintentional movements and has been associated with a number of developmental and clinical conditions, in particular ADHD (Mostofsky et al., 2003; Crasta et al., 2021). Higher total motor overflow scores indicate poorer basic motor control.

Intellectual ability was quantified using the General Ability Index (GAI) derived from the WISC-IV or WISC-V (Wechsler, 2003). We used GAI because we wanted a measure of intellectual ability that was independent of motor control. GAI discounts the impact of tasks involving working memory and processing speed, the latter of which is abnormal in ASD and associated with poor motor control (Mayes and Calhoun, 2008). Higher GAI scores indicated greater intellectual ability.

#### 2.1.3. Study Sample

The study sample for our application of the deconfounded group difference is defined as the subset of participants with a complete set of demographic information (sex, socioeconomic status, and race) and the selected predictors as described in Section 2.3.3. The missingness of the data is depicted in the Web Supplement Figure S1. This subset contains 137 autistic and 348 typically developing children from the original 173 autistic and 373 typically developing children, and we refer to these 485 participants as the complete predictor cases. The socio-demographic characteristics of the complete predictor cases are summarized in Table 1. The impacts of motion exclusion criteria on this subset are discussed in Section 3.1.1.

**Table 1:** Socio-demographic characteristics of complete predictor cases. For continuous variables, mean and standard deviation (SD) are indicated; Kruskal-Wallis rank-sum tests were used to assess diagnosis group differences. For binary and categorical variables, frequencies and percentages are summarized, and differences between diagnosis groups were assessed using either the Chi-square test or Fisher’s exact test. Despite aggregating data from several studies, age and handedness were balanced between diagnosis groups. In contrast, sex, race, and socioeconomic status were imbalanced. ASD=autism spectrum disorder. TD=typically developing. SD=standard deviation.

|  | TD (N=348) | ASD (N=137) | p value |
| --- | --- | --- | --- |
| **Sex** |  |  | 0.002<sup>1</sup> |
| Male | 242 (69.5%) | 114 (83.2%) |  |
| Female | 106 (30.5%) | 23 (16.8%) |  |
| **Age** |  |  | 0.664<sup>2</sup> |
| Mean (SD) | 10.353 (1.249) | 10.286 (1.344) |  |
| Range | 8.020 - 12.980 | 8.010 - 12.990 |  |
| **Race** |  |  | 0.005<sup>3</sup> |
| African American | 36 (10.3%) | 7 (5.1%) |  |
| Asian | 27 (7.8%) | 3 (2.2%) |  |
| Biracial | 45 (12.9%) | 12 (8.8%) |  |
| Caucasian | 240 (69.0%) | 115 (83.9%) |  |
| **Socioeconomic Status** |  |  | 0.007<sup>2</sup> |
| Mean (SD) | 54.072 (9.408) | 51.883 (9.356) |  |
| Range | 18.500 - 66.000 | 27.000 - 66.000 |  |
| **Handedness** |  |  | 0.308<sup>1</sup> |
| Right | 312 (89.7%) | 116 (84.7%) |  |
| Left | 17 (4.9%) | 10 (7.3%) |  |
| Mixed | 19 (5.5%) | 11 (8.0%) |  |
| **Currently On Stimulants** |  |  |  |
| No | 348 (100.0%) | 89 (65.0%) |  |
| Yes | 0 (0.0%) | 48 (35.0%) |  |

<sup>1</sup> Pearson Chi-Square test
<sup>2</sup> Kruskal-Wallis rank sum test
<sup>3</sup> Fisher’s exact test

#### 2.1.4. rs-fMRI Acquisition and Preprocessing

All participants completed at least one mock scan training session to habituate to the MRI environment during a study visit prior to their MRI session. Rs-fMRI scans were acquired on a Phillips 3T scanner using an 8-channel or a 32-channel head coil and a single-shot, partially parallel, gradient-recalled echo planar sequence with sensitivity encoding (repetition time [TR]/echo time = 2500/30 ms, flip angle = 70^◦^, sensitivity encoding acceleration factor of 2, 3-mm axial slices with no slice gap, in-plane resolution of 3.05 × 3.15 mm [84 × 81 acquisition matrix]). An ascending slice order was used, and the first 10 seconds were discarded at the time of acquisition to allow for magnetization stabilization. The duration of rs-fMRI scans varied between 5 min 20 seconds (128 timepoints) and 6.75 min (162 timepoints), depending on the date of enrollment.

Rs-fMRI scans were either aborted or not attempted for seven participants after two unsuccessful mock scan training sessions in the complete predictor case set (3 ASD) due to noncompliance. Rs-fMRI scans for the remaining 478 participants in the complete predictor case set were visually inspected for artifacts and preprocessed using SPM12 (Wellcome Trust Centre for Neuroimaging, London, United Kingdom) and custom code written in MATLAB (The Mathworks, Inc., Natick Massahusetts), which is publicly available (https://github.com/KKI-CNIR/CNIR-fmri_preproc_toolbox). Rs-fMRI scans were slice-time adjusted using the slice acquired at the middle of the TR as a reference, and head motion was estimated using rigid body realignment. Framewise displacement was calculated from these realignment parameters (Power et al., 2012). The volume collected in the middle of the scan was spatially normalized using the Montreal Neurological Institute (MNI) EPI template with 2-mm isotropic resolution (Calhoun et al., 2017). The estimated rigid body and nonlinear spatial transformations were applied to the functional data in one step. Each rs-fMRI scan was linearly detrended on a voxel-wise basis to remove gradual trends in the data. Rs-fMRI data were spatially smoothed using a 6-mm FWHM Gaussian kernel.

#### 2.1.5. Motion QC

We considered two levels of gross motion exclusion:

1. In the lenient case, scans were excluded/deemed unusable if the participant had less than 5 minutes of continuous data after removing frames in which the participant moved more than the nominal size of a voxel between any two frames (3 mm) or their head rotated 3^◦^, where a 3^◦^ rotation corresponds to an arc length equal to 2.6 mm assuming a brain radius of 50 mm (Power et al., 2012) or 4.2 mm assuming 80 mm (Jenkinson et al., 2002; Yan et al., 2013). This procedure was modeled after common head motion exclusion criteria for task fMRI data, which rely on voxel size to determine thresholds for unacceptable motion (Johnstone et al., 2006; Fassbender et al., 2017).
2. In the strict case, scans were excluded if mean FD exceeded .2 mm or they included less than five minutes of data free from frames with FD exceeding .25 mm (Ciric et al., 2017).

Eighty-three participants in the complete predictor case set (17 ASD) completed more than one rs-fMRI scan. For these participants, if more than one scan passed the lenient level of motion QC, we selected the scan with the lowest mean FD to include in our analyses.

#### 2.1.6. Group ICA and Partial Correlations

Thirty components were estimated using group independent component analysis (Group ICA) with 85 principal components retained in the initial subject-level dimension reduction step from the scans that passed lenient motion QC (GIFT v3.0b: https://trendscenter. org/software/gift/; Medical Image Analysis Lab, Albuquerque, New Mexico) (Calhoun et al., 2001; Erhardt et al., 2011). Detailed methods for Group ICA can be found in Allen et al. (2011). We used the back-reconstructed subject-level timecourses for each independent component to construct subject-specific partial correlation matrices (30x30) using ridge regression (*ρ* = 1) (Lombardo et al., 2019; Mejia et al., 2018). After Fisher z-transforming the partial correlation matrices, we extracted the lower triangle for statistical analysis. Following the taxonomy for macro-scale functional brain networks in Uddin et al. (2019), we identified 18 signal components from the 30 group components. A partial correlation is equal to zero if the two components are conditionally independent given the other components. By using the partial correlations, we control for correlations due to the twelve non-signal components, which include some motion artifacts, as well as components mainly composed of white matter or cerebrospinal fluid, which capture other signals of non-interest that impact the brain globally (Bijsterbosch et al., 2020).

### 2.2. Impact of motion QC on the sample size and composition

#### 2.2.1. Impact of motion QC on group sample size

For each level of motion exclusion, Pearson’s chi-squared tests were used to assess whether the proportion of excluded children differed between the ASD and typically developing groups.

#### 2.2.2. rs-fMRI exclusion probability as a function of phenotypes

We used univariate generalized additive models (GAMs) to examine the relationship between the log odds of exclusion and seven covariates: ADOS (ASD group), SRS, inattention, hyperactivity/impulsivity, motor overflow, age, and GAI. We used the subset of children included in the final study sample (Section 2.1.3) for the strict and lenient motion exclusion criteria. We used automatic smoothing determined using random effects with restricted maximum likelihood estimation (REML) (Wood, 2017). We used univariate models rather than a model with all covariates simultaneously because some of the variables are correlated, such that the impact of each variable on rs-fMRI usability may be difficult to estimate. These models are related to the propensity models that will be used in the estimation of the deconfounded group difference (Section 2.3.1). We did not include SES, sex, or race in these models because these variables are not included in the propensity model in Section 2.3.3 due to imbalance between diagnosis groups. Results when controlling for these variables were highly similar (not shown). While the propensity models use an ensemble of machine learning models to predict usability from multiple predictors, our focus for this analysis is on interpretable models. We controlled for multiple comparisons using the false discovery rate (FDR) for the seven univariate models, in which FDR is applied separately to the lenient and strict criteria models (Benjamini and Hochberg, 1995). Although FDR correction was popularized by high-throughput studies conducted in computational biology, Benjamini and Hochberg (1995) originally illustrated the utility of their approach for controlling the expected number of falsely rejected null hypotheses using a study in which a moderate number of tests (15) were performed, which is comparable to our analysis.

We also conducted a similar analysis using univariate GAMs assuming Gaussian errors to examine how the phenotypes are related to mean FD. We conducted separate analyses for the study sample (both usable and unusable cases), children passing the lenient criteria, and children passing the strict criteria, again using FDR correction for seven comparisons within each sample. We also examined whether mean FD differed by sex for these three samples using Mann-Whitney U-tests.

#### 2.2.3. Impact of motion QC on distributions of phenotypes among children with usable data

We examined how the distribution of ADOS (ASD group), SRS, inattention, hyperactivity/impulsivity, motor overflow, age, and GAI differed between included and excluded participants. For additional insight into how scan exclusion may differentially affect autistic versus typically developing children, we stratified this analysis by diagnosis. We visualized the densities using kernel density estimation with default bandwidths in ggplot2 (Wickham, 2016). We then used one-sided Mann-Whitney U tests to test for differences between included and excluded participants for each measure stratified by diagnosis. We also calculated effect sizes as *Z/√N* . We hypothesized that 1) included children would have less severe social, inattentive, hyperactive/impulsive, and motor deficits than excluded children, and 2) included children would be older and have higher GAI. We controlled for multiple comparisons by applying the FDR separately to the thirteen tests (7 for the ASD group and 6 for the typically developing group) performed for the lenient and strict motion QC cases.

#### 2.2.4. Functional connectivity as a function of phenotypes

We also characterized the relationship between phenotypes and functional connectivity. For each level of motion exclusion, we used univariate GAMs to examine the relationship between each phenotypic measure and the adjusted residuals for each edge of signal-to-signal components in the partial correlation matrix. The adjusted residuals are the same data inputted to the deconfounded group difference and are calculated from the residuals of a linear model with mean FD, max FD, the number of frames with FD*<* 0.25 mm, sex, race, socioeconomic status, and diagnosis with the effect of diagnosis added back in as described in Section 2.3.3. Smoothing was determined using the random effects formulation of spline coefficients with restricted maximum likelihood estimation (REML) (Wood, 2017).

#### 2.3. Addressing data loss and reducing sampling bias using the deconfounded group difference

#### 2.3.1. Theory: Deconfounded group difference

Our goal is to estimate the difference in average functional connectivity between autistic children without an intellectual disability and typically developing children. We will use a causally informed approach to correct this associational estimand for potential selection bias following exclusion due to failed motion QC. Let *Y* be a random variable denoting the functional connectivity between two locations (or nodes defined using independent component analysis) in the brain. In practice, these will be indexed by *v* and *v*^′^, but we suppress this notation for conciseness. Let *A* denote the diagnosis indicator variable equal to one if the participant has ASD and zero otherwise. We first consider the hypothetical case in which all participants have usable rs-fMRI data. We use the potential outcomes notation and let *Y* (1) denote the functional connectivity in this hypothetical world (Hernan and Robins, 2020). Our counterfactual *Y* (1) is not functional connectivity if assigned a diagnosis of autism, but rather functional connectivity under an intervention that reduces motion to an acceptable level in all children. Let *W* denote the covariates, which include measures that may be related to functional connectivity and ASD severity. Our *parameter of interest* is the difference in functional connectivity between autistic and typically developing children: *ψ*^*^ = *E*^*^(*Y* (1)|*A* = 1) − *E*^*^(*Y* (1)|*A* = 0), where *E*^*^() denotes an expectation with respect to the probability measure of {*Y* (1)*, A, W* }. It is important to observe that *W* is not independent of *A*, as the distribution of some behavioral variables differ by diagnosis group. We rewrite *ψ*^*^ using the law of iterated expectations to gain insight into our parameter of interest:

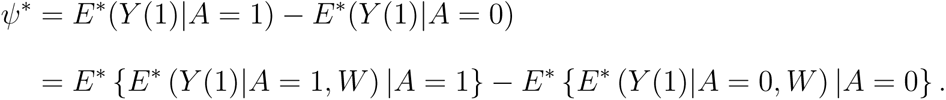

Here, the outer expectation integrates across the conditional distribution of the variables given diagnosis. This associational estimand differs from an average treatment effect (ATE) commonly considered in causal inference (Hernan and Robins, 2020), which integrates across the distribution of the covariates for the pooled population (autistic and typically developing children) to contrast the counterfactual of being assigned a diagnosis of autism with being assigned typically developing.

In contrast to the hypothetical world, many children in the observed world move too much during their rs-fMRI scan for their data to be usable, but we are still able to collect important behavioral and socio-demographic covariates from them. We regard data that fail motion quality control as “missing data.” Let Δ denote a binary random variable capturing the missing data mechanism that is equal to one if the data are usable and zero otherwise. Then data are realizations of the random vector {*Y, A, W,* Δ}. Expectation with respect to their probability measure is denoted *E*() (no asterisk). Additionally, *Y* |(Δ = 0) is missing. Then the naïve difference is *ψ_naive_* = *E*(*Y* |Δ = 1*, A* = 1) − *E*(*Y* |Δ = 1*, A* = 0). We define confounding as *ψ*^*^ ≠ *ψ_naive_* (Greenland et al., 1999). Bias can occur when a covariate is related to data usability/missingness, *W* ↔ Δ, and also related to functional connectivity, *W* ↔ *Y* . Then if the covariate is related to diagnosis, i.e., *W* ↔ *A*, we have *ψ*\* ≠ *^ψ^_naive_*. If there are interactions between *W* and *A*, then we can also have *ψ*^*^ ≠ *ψ_naive_*. These relationships are summarized in the graph in Figure 1. We now define our *target parameter* as a function of usable data. We call this quantity the *deconfounded group difference*:

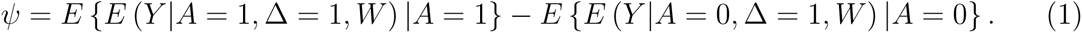

The mathematical distinction between this and the naïve estimator is that in the naïve estimator, *E*(*Y* |Δ = 1*, A* = 1) = *E* {*E*(*Y* |Δ = 1*, A* = 1*, W*)|Δ = 1*, A* = 1}, which differs from *E* {*E* (*Y* |Δ = 1*, A* = 1*, W*) |*A* = 1}, with a similar distinction for *E*(*Y* |Δ = 1*, A* = 0). In the deconfounded group difference, we integrate across the conditional distribution of phenotypic variables given diagnosis versus the naïve approach that integrates across the conditional distribution of phenotypic variables given diagnosis and data usability. We will show in Section 3.1.3 that the distribution of phenotypic variables given diagnosis differs from the distribution given diagnosis and data usabilty.

Identifying the parameter of interest *ψ*^*^ from the target parameter *ψ* requires three assumptions:

(A1.1) *Mean exchangeability*:

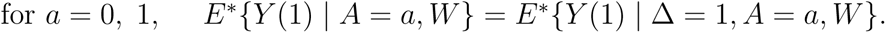

(A1.2) *Positivity*: for *a* = 0, 1 and all possible *w*, *P* (Δ = 1 | *A* = *a, W* = *w*) *>* 0.

(A1.3) *Causal Consistency*: for all *i* such that Δ*_i_* = 1, *Y_i_*(1) = *Y_i_*.

Assumption (A1.1) implies that *W* is sufficiently rich as to contain all variables simultaneously associated with mean functional connectivity and exclusion due to failed motion QC. This assumption is also called ignorability or the assumption of no unmeasured confounders. In the missing data literature, this is closely related to the assumption that data are missing at random: *P* (Δ = 1|*Y, A* = *a, W*) = *P* (Δ = 1|*A* = *a, W*) (van der Laan and Robins, 2003). Assumption (A1.2) implies that there are no phenotypes in the population who uniformly fail motion QC. Assumption (A1.3) stipulates that *Y* from children with usable fMRI data is the same as the outcome that would have been observed under a hypothetical intervention that allows the child to pass motion control (Vander Weele, 2009). Under A1.1 and A1.3, we have *E*^*^{*Y* (1) | *A* = *a, W* } = *E*{*Y* | Δ = 1*, A* = *a, W* }, which allows us to identify the potential outcomes from the observable data.

We estimate our target using doubly robust targeted minimum loss based estimation [DRTMLE, (Benkeser et al., 2017; van der Laan and Rose, 2011)], which involves three steps enumerated below and illustrated in Figure 1 .

1. Fit the propensity model: *P* (Δ|*A, W*). This model characterizes the probability that the rs-fMRI data pass motion quality control. It uses all data to fit the model. Then the usable functional connectivity will be weighted by their inverse probabilities of usability (propensities) during step three.
2. Fit the outcome model: *E*(*Y* |Δ = 1*, A, W*). This step estimates functional connectivity from the covariates for participants with usable rs-fMRI data. It then predicts functional connectivity for both usable and unusable participants.
3. Use DRTMLE to combine functional connectivity from the usable subjects weighted by the inverse probability of usability from step 1 with predictions of functional connectivity for all subjects (usable and unusable) from step 2. Here, DRTMLE is applied separately to each diagnosis group, which calculates mean functional connectivity by integrating across the diagnosis-specific distribution of the covariates from usable and non-usable participants.

Steps 1 and 2 use super learner, an ensemble machine learning technique. The super learner fits multiple pre-specified regression models and selects a weight for each model by minimizing cross-validated risk (Polley et al., 2019). Step 3 combines the propensity and outcome models using DRTMLE. An appealing property of DRTMLE is that the estimate of the deconfounded group difference and its variance are statistically consistent even if either the propensity model or the outcome model is inconsistently estimated. See Benkeser et al. (2017). By statistical consistency, we mean that our estimate converges to the true difference as the sample size goes to infinity, which is different from the causal consistency assumption in A1.3. Here, we know that the missingness mechanism is deterministic based on motion, but we are replacing it with a stochastic model that estimates missingness based on the behavioral phenotypes. Details of our implementation on the real data are in Section 2.3.3.

#### 2.3.2. Toy example and tutorial

We simulate a dataset with bias and estimate the deconfounded group difference in a tutorial available at https://github.com/mbnebel/DeconfoundedFMRI/blob/revision/DeconfoundGroupDifference_Tutorial.Rmd. We generate a sample in which approximately 25% of the participants have ASD, which is similar to the real data (approximately 30% ASD). Then we generate a covariate representing ASD severity, denoted *W_c_*, equal to zero for the typically developing children and generated from a log normal distribution in the ASD group (log *µ*=2, sd=0.4). We generate nine additional standard normal variables unrelated to diagnosis. Then for the propensity model, data usability is generated from a logistic regression model, logit(*E*[Δ = 1|*W_c_* = *w_c_*]) = 2 − 0.2 * *w_c_*. At the mean *W_c_* in the simulated ASD group, the effect is −0.2 * 7.4 = −1.5. In the real data, there were non-linearities with steeper slopes at higher ADOS. The slope was approximately -0.077 at the mean ADOS=14.3, leading to −0.077 * 14.3 = −1.1 (see Section 3.1.2). The simulation design resulted in approximately 88% and 60% usable data in the typically developing and ASD groups, respectively, compared to 84% and 72% using the lenient criteria in the real data.

Next, we defined the outcome model using a linear model in which the slope is -0.2 for *W_c_*, 0 for the nine other covariates, and 0 and 1.4 for the typically developing and ASD intercepts, respectively. This simulation design resulted in the correlation between *W_c_* and *Y* equal to -0.56, and the “true” functional connectivity, i.e., *E*^*^[*Y* (1)|*A* = *a*], equal to approximately -0.20 in the ASD and 0 in typically developing groups leading to a between groups Cohen’s d=0.51. In our dataset, ADOS is weakly correlated with the partial correlations (min=-0.21, max=0.17 across 153 edges for children with usable data under lenient motion QC), and the largest naïve Cohen’s d=0.50 (the challenges of calculating a Cohen’s d with DRTMLE are discussed in Section 4.3). We note that a study on sleeping autistic toddlers found correlations between functional connectivity and ADOS as great as -0.78 in certain subgroups and some large effect sizes (*>*0.8) (Lombardo et al., 2019). We generate a random sample equal to 550, then estimate the deconfounded group difference, as depicted in Fig. 2. We will see that in the real data analysis using lenient motion QC, the naïve difference and deconfounded group difference are more similar than this toy example. However, the toy example illustrates the impact of selection bias under a plausible experimental setup.

**Figure 2:**
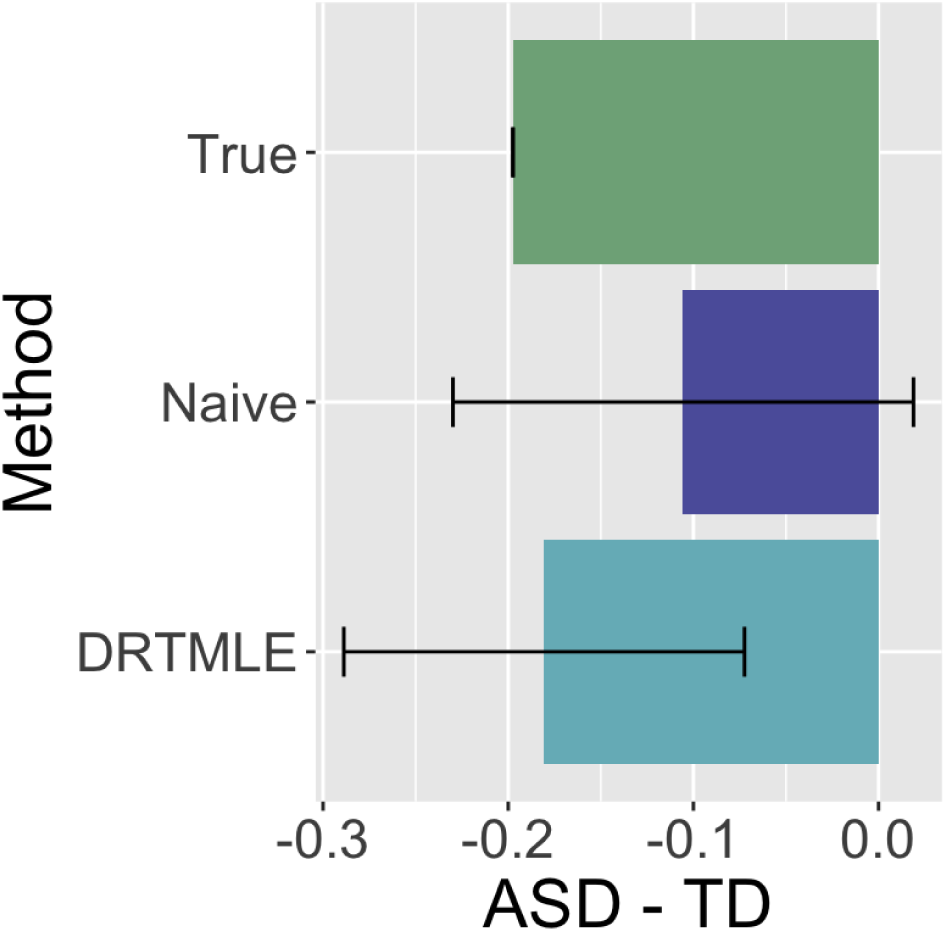
An illustration of the improvement in functional connectivity from DRTMLE compared to the naive approach from a single simulated dataset. The true mean ASD-TD difference in functional connectivity is negative (green bar), with the true mean in the ASD group being negative and the the true mean in the TD group being slightly positive. The estimate of the mean ASD-TD difference from the naïve approach (purple bar) is also negative but closer to zero. Additionally, the 95% confidence interval includes zero. Using DRTMLE, the deconfounded group difference (aqua bar) is closer to the truth and the 95% confidence interval does not include zero. Code to reproduce this example is available at https://github.com/mbnebel/DeconfoundedFMRI/blob/revision/DeconfoundGroupDifference_Tutorial.Rmd.

#### 2.3.3. Application: Deconfounded group difference in the KKI dataset

Recall Step 1 involves fitting a propensity model and Step 2 involves fitting an outcome model (Section 2.3.1). We use the same predictors in the propensity and outcome models: age at scan, handedness (left, right, mixed), primary diagnosis, secondary diagnosis of ADHD, indicator variable for a current prescription for stimulants (all participants were asked to withhold the use of stimulants the day prior to and on the day of the scan), motor overflow, GAI, DuPaul inattention, DuPaul hyperactivity/impulsivity, and ADOS. The ADOS is only administered to children in the ASD group, since it is usually equal or close to zero in typically developing children. We set ADOS equal to zero for all typically developing children. Social responsiveness score was not included due to missing values in 19.5% of observations.

The study sample for our application of the deconfounded group difference is the subset of participants with a complete set of predictors (Section 2.1.3) as depicted in Web Supplement Figure S1. We discuss the study sample and possible biases arising from patterns of missingness in the predictors in Section 4.2.

We focus on the lenient motion QC case because too few participants have usable data following strict motion QC to accurately estimate the outcome model. Functional connectivity metrics based on partial correlations are less sensitive to motion artifacts than those based on full correlations (Mahadevan et al., 2021), but to guard against lingering impacts of motion on functional connectivity and to account for possible confounding due to sampling design, we adjust the partial correlations as follows. For each edge, we fit a linear model with mean FD, max FD, number of frames with FD*<* 0.25 mm, sex (reference: female), race (reference: African American), socioeconomic status, and primary diagnosis (reference: Autism) as predictors. We include sex, race, and socioeconomic status in this model because they differed between autistic and typically developing children (see Section 3.1.1). We then extracted the residuals and added the estimated intercept and effect of primary diagnosis. This approach controls for mean effects that differ by group (ASD versus typically developing) that ideally would be equal, and adjusted residuals have been described in the context of site harmonization in Fortin et al. (2018). Then the “naïve” approach is comparable to the approach used in Di Martino et al. (2014), who included diagnosis, sex, age, and mean FD in a linear model. See Section 4.5 for additional discussion.

<u>Steps 1 and 2</u> (see Section 2.3.1): We use the following learners and R packages when using super learner: multivariate adaptive regression splines in the R package earth (Milborrow, 2011), lasso in glmnet (Friedman et al., 2010), generalized additive models in gam (Hastie, 2020), generalized linear models in glm, random forests with ranger (Wright and Ziegler, 2017), step-wise regression in step, step-wise regression with interactions, xgboost (Chen and Guestrin, 2016), and the intercept only (mean) model; for the outcome model (continuous response), we additionally used ridge from MASS (Venables and Ripley, 2002) and support vector machines in e1071 (Meyer et al., 2021). Parameters were set to their defaults except for the following: the family was equal to binomial (logistic link) in the propensity model with method set to minimize the negative log likelihood; in the outcome models, the method was set to minimize the squared error loss. Note the outcome model is fit separately for each of the 153 edges, whereas the same propensities are used for all edges. The propensity model is fit using the complete predictor cases. The outcome model is fit using the complete usable cases.

<u>Step 3</u>: DRTMLE is applied to ASD for each edge, then to TD. This step uses both the propensities and the predicted outcomes to result in an estimate of the deconfounded mean for the ASD group, the deconfounded mean for the typically developing group, and their variances. We use the non-parametric regression option for both the reduced-dimension propensity and reduced-dimension outcome regression. A z-statistic is formed from their difference under the assumption of independent groups, which is used to test the null hypothesis that functional connectivity is equal in autistic and typically developing children.

Since super learner uses cross validation, its results differ for different random seeds. We ran the entire procedure (propensity model and 153 outcome models) for two hundred different seeds, calculated the DRTMLE-based z-statistic for the difference in functional connectivity, and averaged the z-statistics at each edge from the two hundred seeds. We calculated adjusted p-values using FDR=0.2, which means that we expect 20% of the rejected null hypotheses to be falsely rejected. This threshold has been used in recent papers on FDR (Barber and Cand’es, 2015). We also report edges that survive the more stringent FDR=0.05. We repeated this entire procedure a second time with a different set of 200 seeds. The correlation between the average z-statistics across the 153 edges was greater than 0.99. The same edges were selected at false discovery rate FDR=0.20 in the first and second set of seeds. For the final input to the figures, we pooled both sets of seeds and averaged their z-statistics.

For the naïve approach, we calculated the z-statistic of the average group differences between autistic and typically developing children from the complete usable cases for each of the 153 edges. This test statistic is nearly equivalent to the t-statistic from the linear model with motion variables, sex, socioeconomic status, and diagnosis.

### 2.4. Data and code availability

All data used for this study can be made available by written request through the study’s corresponding author under the guidance of a formal data-sharing agreement between institutions that includes the following: 1) using the data only for research purposes and not attempting to identify any participant; 2) limiting analyses to those described in both institutions IRB-approved protocols; and 3) no redistribution of any shared data without a data sharing agreement.

The code for recreating all analyses, tables, and figures in this study is available at https://github.com/mbnebel/DeconfoundedFMRI/tree/revision.

## 3. Results

### 3.1. Impact of motion QC on the study sample and sample bias

#### 3.1.1. The impact of motion QC on sample size can be dramatic and differs by diagnosis group

Figure 3 illustrates the inclusion criteria used for our analyses and the number of participants remaining after each exclusion step. Missing covariate data excluded 60 participants, or 11% of the total number of participants scanned. Lenient motion QC excluded 19.6% of complete predictor cases, while strict motion QC excluded 66.0% of complete predictor cases. In addition, we found the proportion of excluded children differed by diagnosis group using both levels of motion QC (Figure 3b). Using lenient motion QC, 16.1% of typically developing children were excluded, compared to 28.5% of children in the ASD group (*χ*^2^=8.8, df = 1, p=0.003). Using strict motion QC, 60.1% of typically developing children were excluded, compared to 81.0% of children in the ASD group (p=0.003). Thus, commonly used motion QC procedures resulted in large data losses that more severely impacted the size of the ASD group.

**Figure 3:**
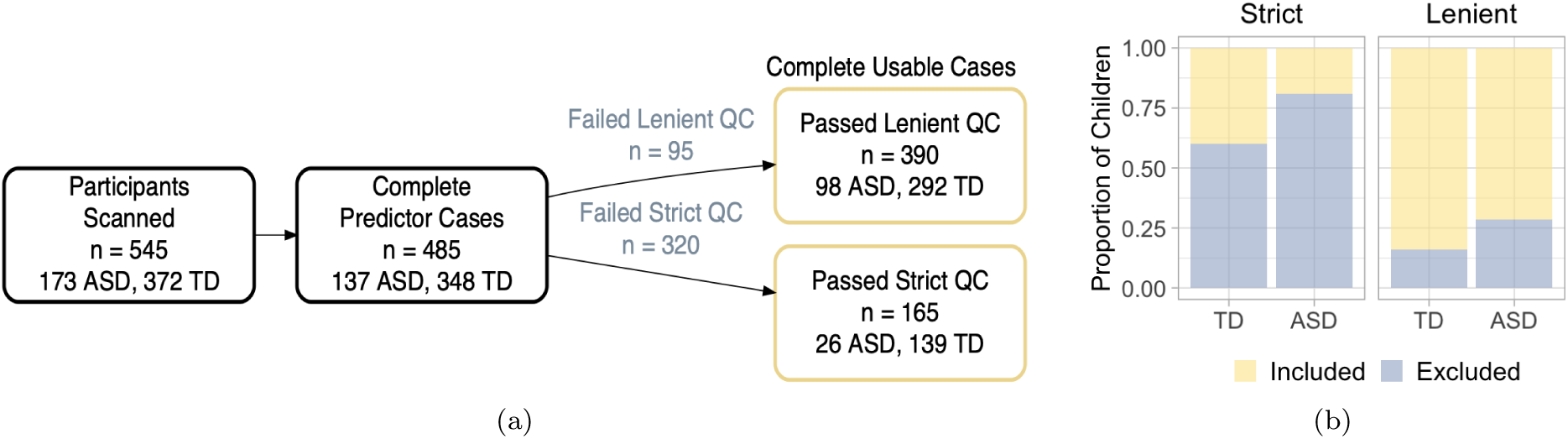
Motion quality control leads to dramatic reductions in sample size. a) Flow chart of inclusion criteria for this study showing the number of participants remaining after each exclusion step. Lenient motion quality control (QC) excluded 19.6% of complete predictor cases, while strict motion QC excluded 66% of complete predictor cases. b) The proportion of children in each diagnosis group whose scans were included (yellow) and excluded (slate blue) using the strict (left) and lenient (right panel) gross motion QC. A larger proportion of children in the autism spectrum disorder (ASD) group were excluded compared to typically developing (TD) children using lenient motion QC (*χ*^2^=8.8, df = 1, p=0.003) and strict (p=0.003).

### 3.1.2. rs-fMRI exclusion probability changes with phenotype and age

We observed that children with higher ADOS scores, SRS scores, inattentive symptoms, hyperactive/impulsive symptoms, or poorer motor control were more likely to be excluded, while older children and children with higher GAI were less likely to be excluded when the lenient motion QC was used (all FDR-adjusted p*<*0.01) as well as the strict motion QC (all FDR-adjusted p*<*0.03) (Figure 4). In particular, there is a sharp increase in exclusion probability using the lenient motion QC for children with higher ADOS scores (slate blue line, left-most panel). The bottom panel of Figure 4 illustrates the covariate distribution for each diagnosis group (pooling included and excluded participants). Interestingly, using the lenient motion QC, the relationship between SRS and exclusion appears flatter over the range of values in the typically developing group and steeper over the range of values in the ASD group (slate blue line). In contrast, the relationship between hyperactivity/impulsivity and exclusion appears linear over the range of values present in the typically developing group but fairly flat over the range of values in the ASD group.

**Figure 4:**
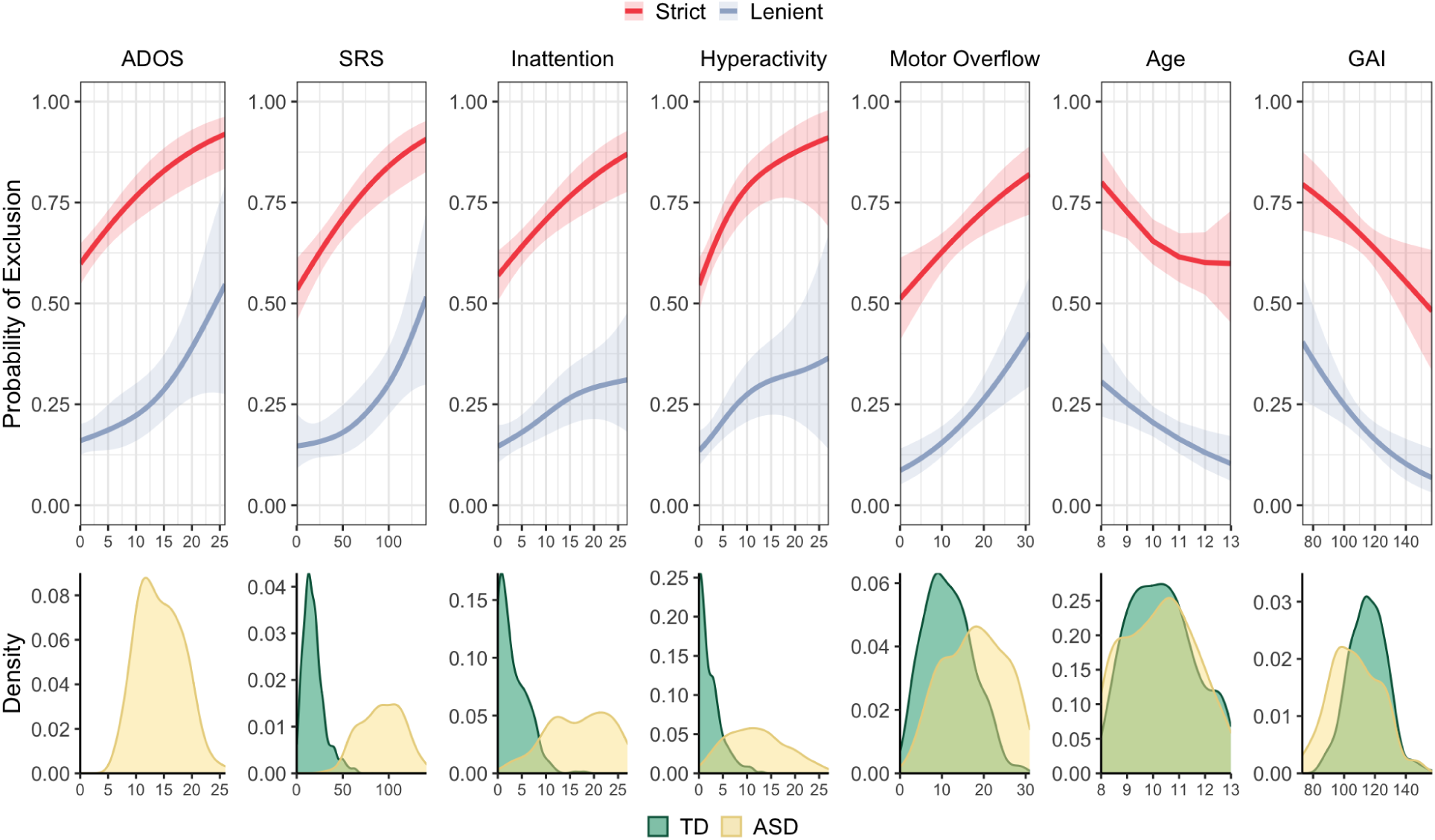
rs-fMRI exclusion probability changes with phenotype and age. Univariate analysis of rs-fMRI exclusion probability as a function of participant characteristics. From left to right: Autism Diagnostic Observation Schedule (ADOS) total scores, social responsiveness scale (SRS) scores, inattentive symptoms, hyperactive/impulsive symptoms, total motor overflow, age, and general ability index (GAI) using the lenient (slate blue lines, all FDR-adjusted p*<*0.01), and strict (red lines) motion quality control (all FDR-adjusted p*<*0.03). Variable distributions for each diagnosis group (included and excluded scans) are displayed across the bottom panel (TD=typically developing, green; ASD=autism spectrum disorder, yellow).

When we considered children with usable and unusable data, we observed similar patterns between mean FD and the covariates as we saw between the probability of exclusion and the covariates. Children with higher (more severe) ADOS scores, SRS scores, inattentive symptoms, hyperactive/impulsive symptoms, or poorer motor control moved more, while older children and children with higher GAI moved less (Web Supplement Figure S2, black lines, all FDR-adjusted p*<*.005). We saw similar patterns following lenient motion QC except that the tendency for children with higher GAI to move more was not significant (FDR-adjusted p=.496). The strict motion QC appears to eliminate the relationship between motion and behavioral phenotypes (FDR-adjusted p*>* 0.05). However, this may be due in part to restricting the distribution of phenotypes, as highlighted by the change in the distribution of ADOS. These biases are further examined in the next section. Before scan exclusion, there was a tendency for boys to move more than girls (p=0.03, uncorrected, r=0.10), but there was not a significant difference between boys and girls with usable data in the lenient (p=0.27) or strict samples (p=0.17).

#### 3.1.3. Phenotype representations differ between included and excluded children

Figure 5 illustrates distributions of the covariates for included and excluded participants stratified by diagnosis group and motion QC level. For the lenient motion QC, median values for included and excluded participants, effect sizes, and FDR-adjusted p values for each measure and diagnosis group are summarized in Web Supplement Table S1. Using the lenient motion QC, we observed biases in both the ASD and typically developing groups toward the selection of older children (FDR-adjusted p=0.08, 0.05 for the ASD and typically developing groups, respectively) with higher GAI (FDR-adjusted p=0.07 for both diagnosis groups). In the ASD group, we also observed biases toward the selection of children who had lower total ADOS, SRS, or motor overflow scores (FDR-adjusted p=0.07, 0.05, and 0.01, respectively), but we did not observe differences in terms of inattentive or hyperactive/impulsive symptoms between included and excluded participants (FDR-adjusted p*>*0.6 for both covariates). In the typically developing group, we did not observe a bias in terms of SRS or inattention (FDR-adjusted p=0.5, 0.3), while there was some evidence of bias for motor overflow (p=0.08). We did observe a bias towards the selection of typically developing children with lower hyperactive/impulsive scores (FDR-adjusted p=0.07). Significant p-values were associated with small effect sizes (*r* = 0.08 to 0.27).

**Figure 5:**
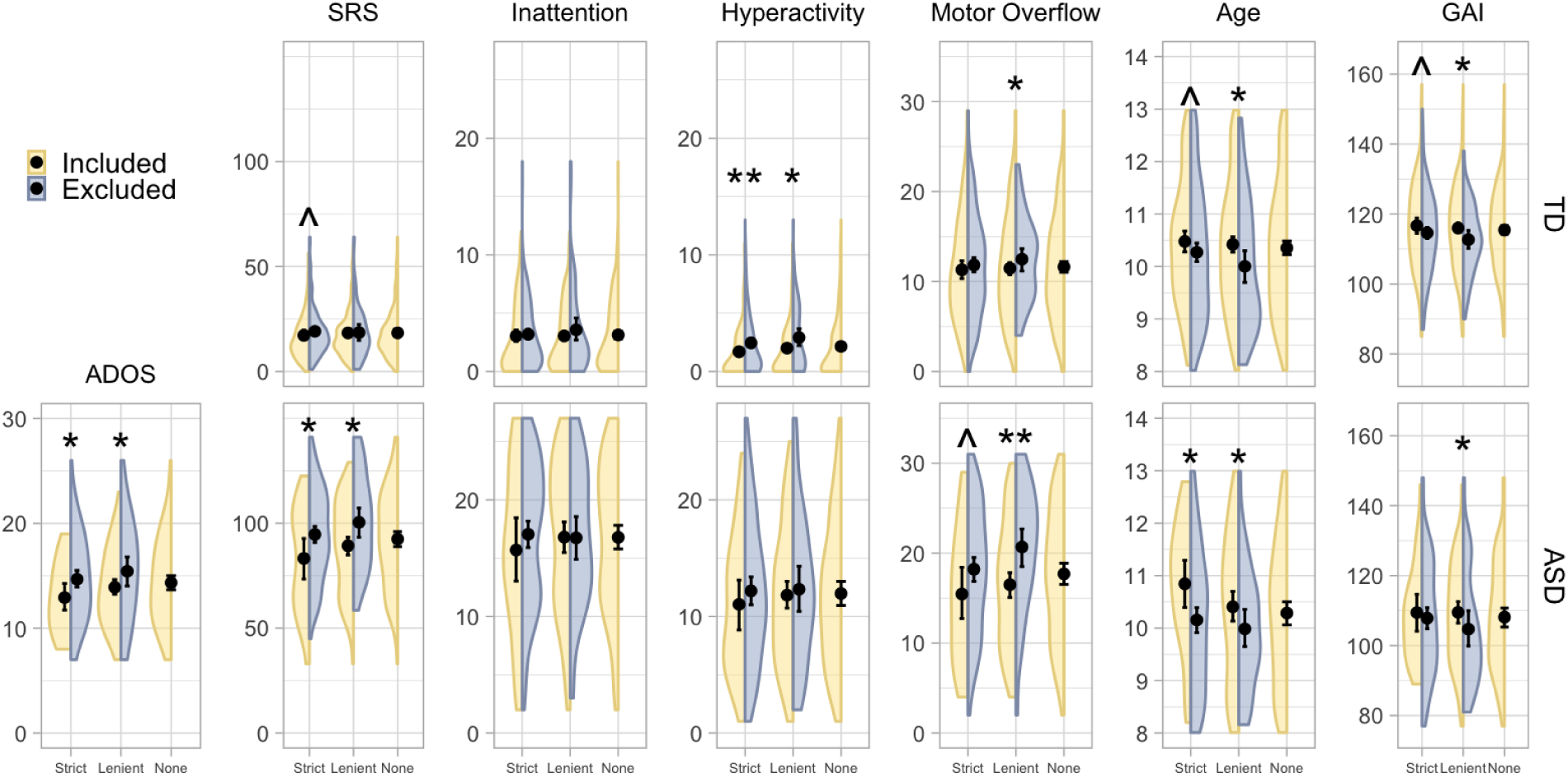
Participants with usable rs-fMRI data differed from participants with unusable rs-fMRI data. Comparison of Autism Diagnostic Observation Schedule (ADOS) scores, social responsiveness scale (SRS) scores, inattentive symptoms, hyperactive/impulsive symptoms, motor overflow, age, and general ability index (GAI) for included (yellow) and excluded (slate blue) participants stratified by diagnosis group and motion exclusion level. The deconfounded mean integrates across the diagnosis-specific distribution of usable and unusable covariates for the variables described in Section 2.3.3, which here is labeled as “None.” Mean values are indicated by a black dot; 95% bootstrap confidence intervals are indicated with black bars. We controlled for 13 comparisons performed for the lenient and strict motion QC cases using the false discovery rate (FDR). ** indicate differences between included and excluded participants with an FDR-adjusted p value *<*0.05; * indicate FDR-adjusted p values *<*0.1; ^indicate FDR-adjusted p values *<*0.2. A larger number of significant differences are observed using the lenient motion QC than the strict motion QC, but very few participants pass strict motion QC. autism spectrum disorder (ASD), typically developing (TD). The R code to produce these split violin plots was adapted from DeBruine (2018).

Differences between included and excluded children also tended to occur using the strict criteria, although in general significance was reduced, owing in part to the reduced sample size in the included group but also to some reduced effect sizes (i.e., motor overflow). Typically developing children who were included were less hyperactive/impulsive than typically developing children who were excluded (FDR-adjusted p=0.02). Median values for included and excluded participants, effect sizes, and FDR-adjusted p values for each measure and diagnosis group are summarized in Web Supplement Table S2.

#### 3.1.4. Phenotypes are also related to functional connectivity

The relationships we observed between rs-fMRI data usability and the covariates examined in the preceding analyses may impact our parameter of interest if those measures are also related to functional connectivity. Figure 6 illustrates histograms of p values for GAMs of the relationship between edgewise functional connectivity (adjusted for sex, SES, race, and motion, see Section 2.3.3) and ADOS, SRS, inattentive symptoms, hyperactive/impulsive symptoms, total motor overflow, age, and GAI across participants with usable rs-fMRI data using the lenient motion QC (slate blue bins) and the strict motion QC (red bins). This analysis is related to the outcome model used in the deconfounded group difference, as it provides insight into whether the sampling bias will impact the mean difference in functional connectivity between groups. Here, we focus on a single phenotype in each GAM for interpretability. For a given phenotype, a clustering of p values near zero suggests that a covariate is associated with functional connectivity for a greater number of edges. If there is no association between the covariate and functional connectivity, we expect the p values to be more uniformly distributed. We see strong clustering of p values near zero for total ADOS across participants with usable rs-fMRI data under lenient and strict motion QC. For SRS, we see some clustering using participants who pass lenient and strict motion QC. For inattentive symptoms, we see a clustering of p values near zero using participants who pass lenient motion QC. For hyperactive/impulsive symptoms, we see a clustering of p values near zero following lenient and strict motion QC. For motor overflow, we see some clustering of p values near zero for both levels of motion QC. For age, we see a clustering of p values near zero using participants who pass the lenient motion QC but not strict. For GAI, we see a clustering of p values near zero using participants who pass the strict motion QC but not lenient.

**Figure 6:**
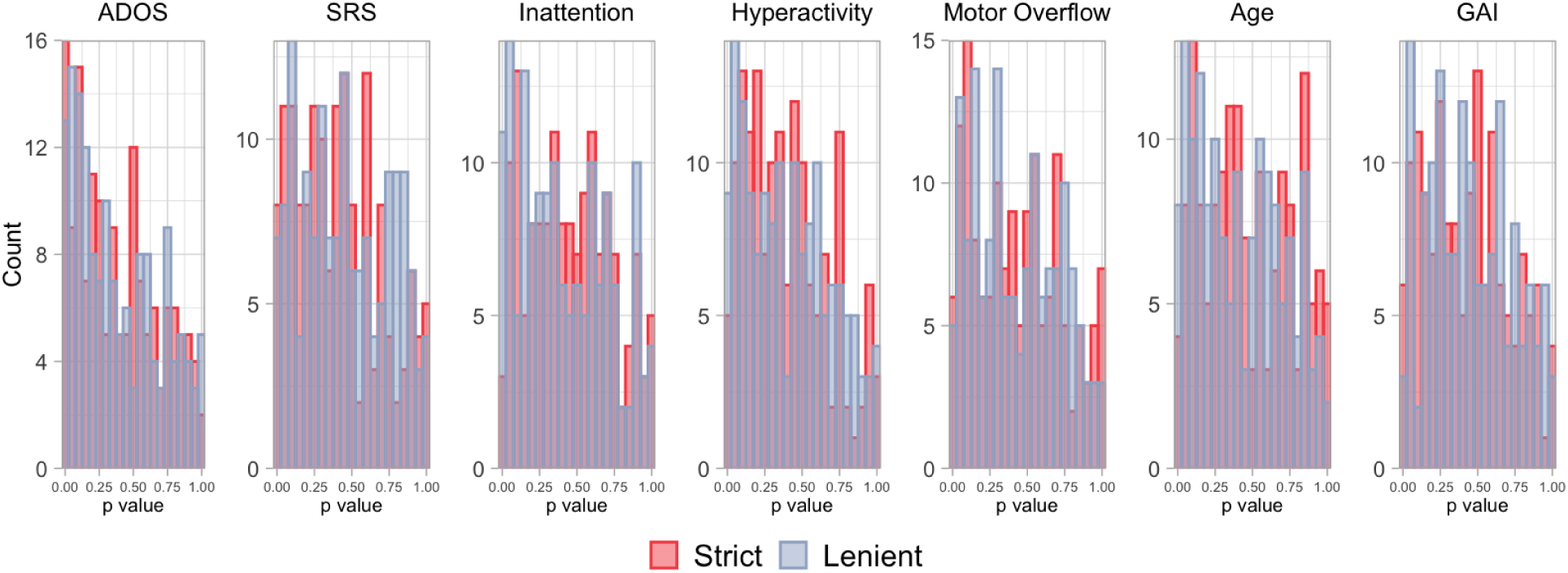
Some covariates related to rs-fMRI exclusion probability are also related to functional connectivity. Histograms of p values for generalized additive models of the relationship between edgewise functional connectivity in participants with usable rs-fMRI data and (from left to right) ADOS, social responsiveness scale (SRS) scores, inattentive symptoms, hyperactive/impulsive symptoms, total motor overflow as assessed during the Physical and Neurological Exam for Subtle Signs, age, and general ability index (GAI). For a given covariate, a clustering of p values near zero suggests that covariate is associated with functional connectivity for a greater number of edges. Several covariates appear to be related to functional connectivity using both the lenient motion quality control (slate blue bins) and the strict motion quality control (red bins).

### 3.2. Application: Deconfounded group difference in the KKI Dataset

We examined the stability of the propensity scores across all random seeds. Propensities near zero can increase the bias and variance of causal effects (Petersen et al., 2010) and indicate a possible violation of the positivity assumption (A1.2). The smallest propensity ranged from 0.23-0.52. This indicates that there is a reasonable probability of data inclusion across the range of {*W, A*} and that Assumption (A1.2) is likely to be adequately satisfied. The AUCs for predicting usability across all seeds ranged from 0.67 to 0.99, and on average was 0.86, whereas the AUC was 0.68 using logistic regression and 0.69 using a logistic additive model, which indicates that the super learner often improves the accuracy of the propensity model.

The deconfounded group difference estimated using DRTMLE revealed more extensive differences between the ASD and typically developing groups than the naïve approach (Fig. 7, Web Supplement Table S3). At FDR=0.20, the naïve approach indicated four edges showing a negative difference in functional connectivity between the ASD and typically developing groups (ASD*<*TD; red lines) and four edges showing a positive difference (ASD*>*TD; blue lines). The DRTMLE approach also indicated these eight edges, with seven having smaller p values relative to those for the naïve approach. The DRTMLE approach also indicated an additional eight edges showing negative group differences and nine additional edges showing a positive group difference. Network nodes that gained edges from the DRTMLE versus the naïve method (FDR=0.2) included the executive control (+10), default mode (+5), so-matomotor (+8), ventral attention (+5), pontomedullary/cerebellar (+3), dorsal attention (+2), and visual (+1) networks. At FDR=0.05 (Web Supplement Figure S3), the naïve approach only indicated one edge showing a negative difference in functional connectivity between the ASD and typically developing groups (primary visual IC-02 to bilateral executive control IC-27). The DRTMLE approach indicated this edge with a smaller p value, while also indicating five additional edges with a total of three of the six edges showing a negative difference in functional connectivity (ASD*<*TD). Network nodes that gained edges from the DRTMLE versus the naïve method (FDR=0.05) included the default mode (+3), pontomedullary/cerebellar (+3), dorsal attention (+2), and executive control (+1) networks.

**Figure 7:**
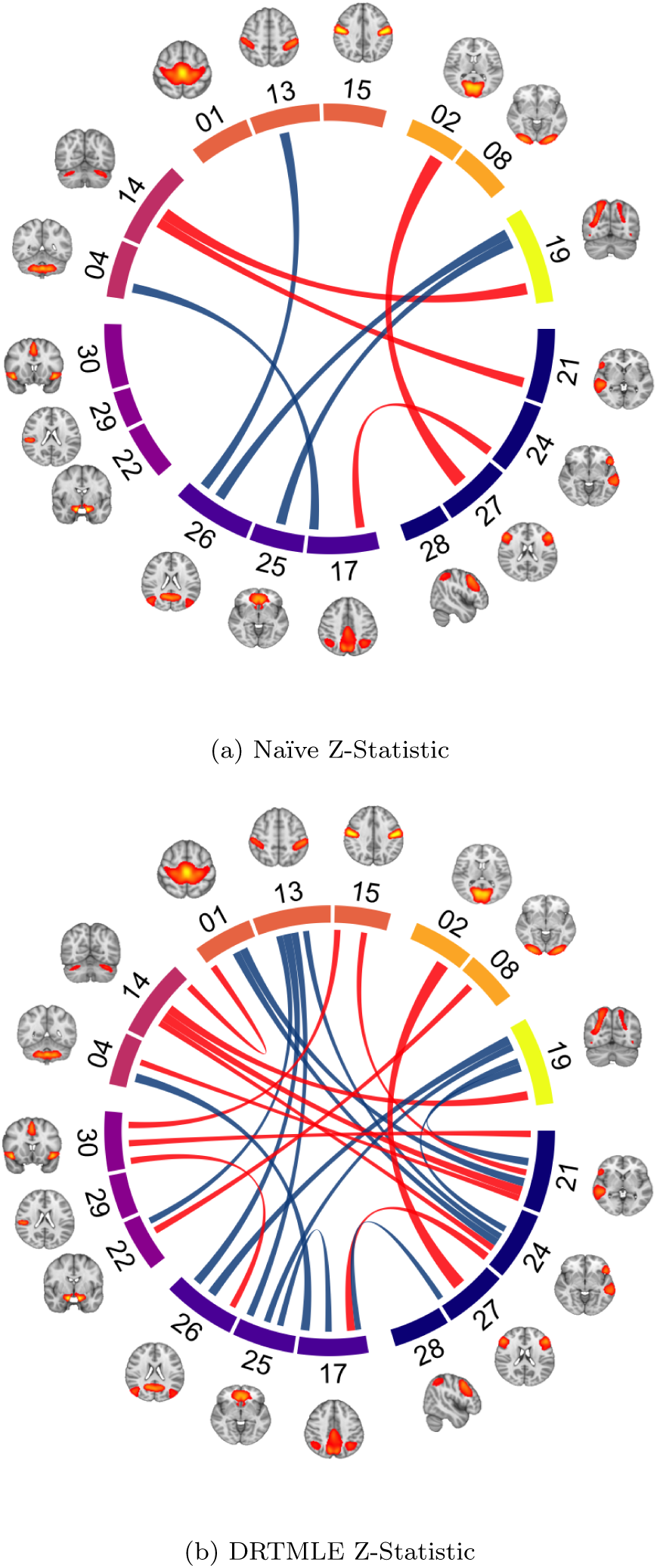
The DRTMLE deconfounded group difference revealed more extensive differences than the naïve approach. Z-statistics for ASD versus TD using a) the naïve test and b) using DRTMLE. Connections are thresholded using a false discovery rate (FDR) of 0.20. Blue lines indicate ASD*>*TD (4 in naïve, 13 in DRTMLE). Red lines indicate ASD*<*TD (4 in naïve, 12 in DRTMLE). Brain regions contributing to each independent component are illustrated and components are grouped by functional assignment. Navy nodes: control. Blue violet: default mode. Purple: salience/ventral attention. Magenta: pontomedullary/cerebellar. Coral: somatomotor. Orange: visual. Yellow: dorsal attention. FDR=0.05 is plotted in Web Supplement Figure S3. See Web Supplement Figure S4 for a visualization of the naïve and deconfounded means and individual-level partial correlations. These plots were generated using the circlize package in R (Gu et al., 2014) and the tutorial provided by Mowinckel (2018).

Functional connectivity scores further from zero reflect stronger functional connectivity regardless of sign; positive scores reflect stronger positive partial correlations, or more integrated intrinsic activity between nodes. Negative scores reflect negative partial correlations, or more segregated intrinsic activity between nodes. The sign of average group effects remained consistent, as did the direction of group differences (Web Supplement Table S3). Edges showing positive group differences in functional connectivity included edges for which positive correlations were strengthened in the ASD group compared to the typically developing group, as well as connections in which negative correlations were weaker in the ASD group compared to the typically developing group. Similarly, the edges showing negative group differences included connections for which negative correlations were strengthened in the ASD group compared to the typically developing group, as well as connections for which positive correlations were weaker in the ASD group compared to the typically developing group.

In this application, the deconfounded means were very similar to the naïve means (Web Supplement Figure S4). Additionally, partial correlations were highly variable, with the range of partial correlations in the ASD and typically developing groups broadly overlapping. However, the small changes in the means also contributed to the more extensive ASD-TD differences in DRTMLE versus the naïve approach depicted in Figure 7. Of the additional 17 edges selected at FDR=0.20 by DRTMLE, eight of the 17 had a decrease in the standard error but all 17 had an increase in the absolute difference between the ASD and typically developing groups.Across all edges, the absolute difference between groups increased in 106/153 edges and the standard error decreased in 81/153. We discuss effect sizes in Section 4.3.

## 4. Discussion

We set out to understand what part of the autism spectrum we are characterizing in rs-fMRI analyses: Does excluding high-motion participants allow us to draw conclusions about average brain function/connectivity that are representative of 8-to-13-year-old children across the entire autism spectrum, or does it introduce bias? The primary message that emerges from our findings is that ignoring bias due to motion exclusion can be problematic scientifically. Using data from a large sample of autistic children without an intellectual disability and typically developing children, we demonstrated that motion exclusion changes the distribution of behavioral and sociodemographic traits in the study sample that are related to functional connectivity. This finding suggests that the generalizability of previous studies reporting naïve analyses may be limited by the selection of older children with less severe clinical profiles because these children are better able to remain still during an rs-fMRI scan. We further propose a statistical approach for addressing the data loss and possible bias following motion QC using DRTMLE. Our findings indicate more extensive differences between autistic and typically developing children using DRTMLE as compared with conventional approaches.

In our study, the impact of motion QC on sample size was dramatic and differed by diagnosis group. Detailed reporting of the number of participants excluded for excessive head motion is far from standard practice, but we found that motion QC removed a larger proportion of autistic children compared to typically developing children, which is consistent with the patterns reported in Redcay et al. (2013) and Jones et al. (2010). Across diagnosis groups, children with more severe social deficits, more inattentive symptoms, more hyperactive/impulsive symptoms, or poorer motor control were more likely to have unusable rs-fMRI data and be excluded, while older children or children with higher intellectual ability were less likely to be excluded for both levels of motion QC. Similarly, Simhal et al. (2021) found that children with ASD and children with ADHD who failed a mock MRI training protocol were younger, had lower verbal and non-verbal intelligence scores, and more severe ADOS scores than children with ASD and children with ADHD who passed the training protocol. These findings suggest that the mechanisms driving missingess in rs-fMRI studies may be related to scientifically relevant participant characteristics.

The estimate of mean functional connectivity should be representative of all children enrolled in the study, assuming the enrolled participants are a representative sample from the target population. However, we observed that participants with usable rs-fMRI data differed from participants with rs-fMRI data that would have been excluded using conventional approaches. Autistic children excluded by lenient motion QC tended to be younger, displayed more severe social deficits (both observed by the experimenter using the ADOS and reported by parents/teachers using the SRS), more motor overflow, or lower intellectual ability than autistic children who were included. We observed similar differences between included and excluded autistic children following strict motion QC, although in general, power was reduced due to the reduced sample size. Moreover, these characteristics are exactly those that showed relationships with functional connectivity among children with usable data following one or both levels of motion QC. The strength of these relationships between clinically relevant measures and functional connectivity among children with usable data appeared to depend on the level of motion QC used. Given that the definition of usability varies widely among rs-fMRI studies, our findings suggest that differences in the representation of symptom severity among children with usable data following motion QC may have partially contributed to discrepancies in the literature regarding ASD-associated functional connectivity findings. To improve comparison across studies, it is critical for rs-fMRI researchers to transparently assess the amount of information lost following motion QC (Fig. 3), to consider whether participant characteristics related to usability are also related to the effect of interest, and to try to address the loss of power and potential bias if they are.

Here, we have advanced this issue using techniques from the missing data and causal inference literature combined with an ensemble of machine learning algorithms. Our framework explicitly treats missingness due to motion QC as a source of bias, and we define a target parameter called the deconfounded group difference, which utilizes the distribution of diagnosis-specific behavioral variables across usable and unusable scans. The general concept of this framework is to recognize that children with usable data are not representative of all enrolled children within each diagnosis group. DRTMLE combines the results of inverse propensity weighting and G-computation, which improves robustness relative to either approach alone. Inverse propensity weighting gives more weight to children with more severe symptoms and usable functional connectivity data because a) they are more likely to be missing and b) functional connectivity is related to symptom severity so we need them to stand in for all children with more severe symptoms who are excluded due to data quality concerns. The outcome model estimates functional connectivity for all children, including those with greater symptom severity, and in this sense accounts for children with unusable data. We use an ensemble of machine learning methods to flexibly model possible non-linear relationships between phenotypic traits and data usability (the propensity model) and between phenotypic traits and functional connectivity (the outcome model). For both the propensity and outcome models, we include a rich collection of variables that we expect to be associated with rs-fMRI usability, functional connectivity, or both. Including variables that contribute to both rs-fMRI usability and functional connectivity represents an opportunity to decrease bias. Including variables that contribute to functional connectivity but not necessarily to rs-fMRI usability represents an opportunity to decrease the variance of our estimate without increasing bias. The propensity and outcome models are then combined using DRTMLE, which results in statistically consistent estimation of the deconfounded group difference and its variances under the assumptions in Section 2.3.1 and discussed in Section 4.4.

### 4.1. Possible scientific insights gained from DRTMLE

The pattern of group differences observed using DRTMLE is consistent with knowledge of DMN-DAN interactions, such that the DMN shows task-induced deactivation, whereas the DAN shows task-induced activation (Padmanabhan et al., 2017). Findings from task-based fMRI studies suggest that individuals with ASD show lower deactivation of the DMN during self-referential processing tasks as compared to typically developing controls (Kennedy et al., 2006; Padmanabhan et al., 2017). Recent findings also suggest a crucial role of the posterolateral cerebellum, a region functionally connected to the DMN (Buckner et al., 2011), in both social mentalizing (Van Overwalle et al., 2020) and behaviors central to a diagnosis of ASD (Lidstone et al., 2021; Stoodley et al., 2017). The cerebellum is also believed to form and update internal models of the world for predictive control in both social and nonsocial contexts (Blakemore et al., 2001). Functional connectivity between the DMN, DAN, and cerebellum should be a focus of future research to better understand the neural mechanisms contributing to autism.

### 4.2. The target population, other possible biases, and methods to address them

Although DRTMLE can address issues of missing data due to motion QC, it does not address possible biases in the sample of children with behavioral data that appear in the study, which may differ from the target population. The target population in the original data-collecting studies was children without an intellectual disability. Within this target population, our study is a convenience sample in that participants meeting the eligibility criteria were recruited using flyers and patient records, rather than randomly sampled. In fact, we had a five-to-one ratio of boys to girls in our ASD sample compared to the estimated four-to-one relative risk nationwide (Maenner et al., 2021). This convenience sampling approach is an important shortcoming in many fMRI studies. A recent study found a healthy volunteer bias in the UK Biobank sample (Fry et al., 2017). Bradley and Nichols (2022) used propensity score weighting methods to decrease this bias in a study of brain structure but noted that these approaches require access to external individual-level population data, which is often unavailable. Extending the DRTMLE approach to accommodate situations in which the study sample deviates from the target population is an important area for future work.

A goal of the original data-collecting studies was to estimate the association between functional connectivity and a diagnosis of ASD separate from any association between functional connectivity and intellectual disability. A recent cohort-based study of heritability and familial risk indicated that ASD without a co-occurring intellectual disability may have a greater genetic basis than ASD with an intellectual disability (Xie et al., 2020), suggesting that these ASD phenotypes may be etiologically distinct. Approximately 35% of autistic children have an intellectual disability (Maenner et al., 2021), and our findings may not generalize to this segment of the autism population (Reiter et al., 2019). Unless we as a field make a concerted effort to correct for the over-representation of cognitively abled children in autism research, we will never be able to answer this question. In addition, an estimated 70% of autistic children have at least one comorbid disorder (Simonoff et al., 2008). The original data-collecting studies allowed for two of the most common comorbid mental health disorders, namely ADHD and social anxiety disorder, but excluded all others. Because research informs the revision of diagnostic criteria and health and educational policy, we must improve recruitment and data collection from children we have historically excluded from rs-fMRI research of autism.

Scanning these underrepresented populations requires thoughtful experimental accommodations. Some approaches for improving the likelihood of collecting usable data from pediatric participants focus on scan preparation: having a caregiver model scan procedures, practicing in an MRI simulator (Nordahl et al., 2016; Horien et al., 2020; Simhal et al., 2021), or playing a virtual reality-based MRI game (Stunden et al., 2021; Pua et al., 2020). Others focus on modifications during the scan: passive movie-viewing to reduce boredom (Vanderwal et al., 2019), providing real-time feedback to participants using framewise integrated real-time MRI monitoring (FIRMM) software (Dosenbach et al., 2017; Greene et al., 2018), or using personalized incentive systems to reward compliance with MRI instructions (Pua et al., 2020). The efficacy of these strategies varies with age, and some strategies come at a cost. For instance, extensive MRI simulator practice has been used to scan small samples of autistic children with intellectual disabilities (Nordahl et al., 2016), but this places additional burdens on families by requiring multiple visits. In addition, some studies with autistic children have observed that head motion, while reduced after extensive training, is still associated with symptom severity (Simhal et al., 2021; Gabrielsen et al., 2018). Virtual reality-based games played at home may be as effective as MRI simulator training for typically developing children, (Stunden et al., 2021); however, it remains unclear if children with neurodevelopmental disorders would respond similarly. Investigating in-scanner strategies, Greene et al. (2018) found that passive movie-viewing and real-time feedback both reduced head motion, but movie-viewing changed functional connectivity. Finn and Bandettini (2021) found that changes in functional connectivity elicited by movie-viewing in adults aided in the prediction of behavioral traits, but similar work has not yet been conducted in children. As we develop methods for increasing participant compliance, our tolerance for what constitutes acceptable levels of head motion will likely change (for instance, Marek et al. (2019) excluded 40% of the ABCD study sample which was collected using FIRMM, while Marek et al. (2022) excluded 60%). We hope our proposed approach will be used in combination with some of these experimental strategies to advance studies including important subgroups of autistic children.

While our method represents an important first step towards addressing bias associated with motion QC, its effectiveness is limited by the missingness of the predictors. Our application assumes that the predictors are missing completely at random. In the case of motor overflow and GAI, missingness may have occurred because some children were unable to complete the cognitive and behavioral tests. The current study focuses on the possible bias due to missingness in the outcome. Imputation to address possible bias due to missingness in the covariates in the context of our application to functional connectivity deserves its own treatment. Multiple imputation involves generating multiple datasets to incorporate uncertainty in the imputation process and involves a variance adjustment for the between-imputation variance (Little and Rubin, 2019). Ideally, all relationships that will be investigated in the analysis should be included in the imputation process, including relationships with the outcome (Azur et al., 2011), as Moons et al. (2006) found imputation that did not include the outcome resulted in large biases. However, our application includes 153 outcomes (each edge) that contain a biased pattern of missingness (the bias we are correcting using DRTMLE), and it is unclear how to best handle this situation.

### 4.3. Similarities between naïve and deconfounded means, sample size limitations, inference, and effect size

When using the lenient criteria on the KKI dataset, the bias corrections using DRTMLE were small (Web Supplement Figure S4). This may be due to weak relationships between the phenotypic variables and the partial correlations, which is consistent with recent ABCD study results suggesting that the largest, replicable associations between functional connectivity and behavioral phenotypes ranged from *r* = 0.14 − 0.34 (Marek et al., 2022). Additionally, our data processing and the use of partial correlations from group ICA may have decreased motion artifacts at the expense of attenuating associations. Our simulated example in Fig. 2 provides an example where the relationship with autism severity is stronger, and we see larger bias.

Studies using stricter criteria may induce larger biases, in which case there may be larger differences between the DRTMLE means and the naïve means. We were unable to apply DRTMLE following strict motion QC because only 29 autistic children had usable data. Visual inspection of the densities in Figure 5 reveals that the sampling bias was larger in the strict versus lenient case for all phenotypes. We speculate that the stricter criteria would result in larger differences between DRTMLE and the naïve approach.

DRTMLE can be used to address data loss by improving efficiency, which can result in smaller standard errors relative to the naïve approach. The TMLE framework leverages all available covariate data, and when the covariate data are predictive of the outcome, this can improve statistical power (Moore and van der Laan, 2009). One potential limitation is that DRTMLE underestimates the variance of group estimates for small sample sizes, resulting in anti-conservative p-values (Benkeser et al., 2017). We cannot disentangle the possible gains in efficiency from the possibly anti-conservative p-values (due to a finite sample). This limitation would be more of a concern following strict motion QC; in that case, only 29 autistic children were labeled as having usable scans. However, using the lenient motion QC, 98 autistic participants and 292 typically developing children had usable scans. In addition, the FDR corrected p-values we use are conservative in the sense that they do not leverage the positive correlations between some edges. An important avenue for future research is to use permutation tests for inference (Winkler et al., 2014) with DRTMLE. Permutation tests can result in finite sample inference while improving power using max statistics, but they create computational challenges.

As in many other rs-fMRI studies, we observed extensive variability among participants in the modified partial correlations used as input to DRTMLE (Web Supplement Figure S4). This variability resulted in generally small effect sizes from the naïve approach. The maximum Cohen’s D across 153 edges was 0.50 at IC02-IC27, which is a medium effect size, and the average naïve effect size among the twenty-five edges selected by DRTMLE at FDR=0.20 was 0.26. Unfortunately, calculating effect sizes in DRTMLE is an open problem. We would need to define a new parameter of interest, the population pooled standard deviation, under the counterfactual that all data are usable and then define an estimator of this parameter. When calculating Cohen’s d in a two sample t-test setting, the standard errors of each mean can be multiplied by ^√^*n*_1_ and ^√^*n*_2_ to recover each standard deviation, which can then be pooled. In DRTMLE, this would not result in an estimate of the standard deviations of each group under the counterfactual of all usable data, since the standard errors are derived from the influence function of *ψ*. This is an important avenue for future research.

Another limitation of the current study is that machine learning algorithms typically require a relatively large sample size compared to classic approaches. We use cross-validation to guard against overfitting, which has been shown to be effective even without having an independent test dataset (Benkeser et al., 2019). One drawback of cross-validation approaches is that they can be sensitive to the random seed. We addressed this limitation by repeating the cross-validation hundreds of times. Each estimation routine takes approximately six hours on a single core (2.60 GHz), which includes fitting the propensity model and 153 edge outcome models. We used a high performance cluster and 100 cores, and conducted two sets of 200 seeds, such that the full estimation routine took approximately 24 hours. The average z-statistic from the two sets were nearly equivalent.

### 4.3. Model assumptions and possible violations

Estimating the difference in functional connectivity between autistic and typically developing children in the counterfactual world in which all data are usable from the observable data involves three assumptions: mean exchangeability, positivity, and consistency of the counterfactual and the observed outcome (causal consistency) (Section 2.3.1).

With respect to mean exchangeability or the assumption of no unmeasured confounders, we assume that functional connectivity is independent of the missingness mechanism given our variables {*W, A*}. As noted, the missingness mechanism is deterministic based on head motion, but we are replacing it with a stochastic model that estimates missingness from {*W, A*}. In our application, it is important that summary measures of head motion were *not* included in the propensity and outcome models. To understand the reason for this, consider that children who nearly fail motion QC may have some motion impacts in their functional connectivity signal. The deconfounded group difference assumes that *Y* reflects the signal of interest, i.e., neural sources of variation that are not corrupted by motion. We took several steps to account for potential motion impacts on functional connectivity in children who nearly fail; we used partial correlations from an ICA that includes some motion artifact components (which removes these sources of variance) and residuals from a linear model including motion, as described in Section 2.1.6 and Section 2.3.3, which results in a *Y* that more closely captures neural sources of variation. However, if we then included summary motion measures in our propensity and outcome models, the propensity model would up-weight these children who nearly failed, and the outcome model, integrating over the full range of head motion, would potentially reintroduce the motion impacts we tried to carefully remove. Additionally, our statistical estimator has the double robustness property: if at least one of the propensity or outcome models is correctly specified, we obtain a statistically consistent estimator of the deconfounded group difference (Bang and Robins, 2005; Benkeser et al., 2017). We include a rich set of predictors and an ensemble of machine learning algorithms, which helps to address the assumption of no unmeasured confounding.

Positivity assumes that there are no values of {*W, A*} such that the data will always be unusable. Violations of positivity assumptions lead to out-of-sample prediction of functional connectivity in the outcome model and instabilities in the propensity model, which can lead to greater variance and bias (Petersen et al., 2010). In Fig. 5, we see that for the lenient criteria, the range of the behavioral traits generally overlap between included and excluded participants, although the most severe ADOS score does not appear among the included children. The highest ADOS score among included children was 23; among all children, 26 (A change in the range also occurs for SRS, but SRS was not included in the propensity and outcome models due to a large proportion of missing values.). As reported in Section 2.3.3, all propensities were greater than 0.30 for the first five random seeds. The lack of propensities close to zero for children with usable or unusable data indicates that the assumption of positivity is reasonable in our application. Regarding the last assumption, causal consistency is a technical assumption that assumes that *Y* (1) is the same as *Y* when a child has usable data, which in general cannot be tested but seems reasonable.

### 4.5. Accounting for variables that should be balanced between diagnosis groups

A possible limitation of the current approach is that we account for covariate imbalance between the ASD and typically developing groups using linear regression prior to attempting to account for bias due to data usability, and it may be desirable to pursue a statistical method that integrates covariate balancing into the deconfounded group difference. The deconfounded group difference estimates the marginal mean of each diagnosis group, where integration is across the distribution of the behavioral variables given diagnosis (defined in Section 2.3.1). However, our typically developing sample was aggregated from multiple rs-fMRI studies conducted at KKI, not all of which involved a comparison sample of autistic children. As a result, sex, race, and socioeconomic status significantly differed between diagnosis groups (Table 1) for this secondary analysis. In an ideal prospective experiment using a random sampling design, these socio-demographic variables would not differ. The naïve approach estimated the difference between autistic and typically developing children while controlling for mean FD, max FD, number of frames with FD*<* 0.25 mm, sex, race, and socioeconomic status in a linear model, which is similar to the approach in Di Martino et al. (2014). Controlling for variables in a linear model corresponds to estimating the conditional mean of functional connectivity given these variables. The residuals of the linear model plus the effect of diagnosis are used as input to estimate the deconfounded group mean for each diagnosis group. If there is no sampling bias due to motion exclusion, then the deconfounded group difference is approximately equivalent to the naïve approach, which is a nice aspect of the present study in that it presents a method for evaluating the extent to which estimates from the conventional approach are biased. Our approach accounts for possible confounding due to the demographic and remaining motion imbalances in the ASD and typically developing samples, although it does so using the traditional linear model.

We can define two sets of variables: 1) variables that we would like to be balanced in autistic and typically developing children in an ideal sample, and 2) variables whose distribution is specific to diagnosis. The target parameter used in estimating an average treatment effect in causal inference marginalizes with respect to the distribution of variables pooled across treatments, which would address biases introduced by the first set of variables. Our deconfounded group difference is associational and addresses the biases introduced by the second set of variables. Future work could define a target parameter that marginalizes with respect to the desired distribution of variables that should be balanced and the desired distribution of variables whose distribution depends on diagnosis.

### 4.6. Other methods to account for bias

We use DRTMLE to estimate the deconfounded group difference. However, a host of other statistical methods could be applied to the same end including covariate matching, propensity score matching (Stuart, 2010; Bridgeford et al., 2021), inverse propensity weighting (Lewinn et al., 2017), G-computation (Robins, 1986; Snowden et al., 2011), augmented inverse propensity weighting (Robins et al., 2012), and targeted maximum likelihood estimation (van der Laan and Rubin, 2006; Schuler and Rose, 2017). Comparing the performance of these approaches in the context of rs-fMRI studies is an important area for future work but is beyond the scope of this paper.

### 4.7. Significance to other neurological disorders and developmental studies

We used an rs-fMRI study of ASD to illustrate the unintended cost of motion QC on study generalizability, but the issue of data loss and selection bias due to motion QC is neither specific to ASD or to rs-fMRI. Head motion-induced artifacts are a notorious problem for all magnetic resonance-based neuroimaging modalities, and the relationship between motion and participant characteristics is problematic in studies of developmental and aging trajectories, as well as other neurological disorders. For instance, we found that younger children were more likely to be excluded. Recent studies investigating associations between functional brain organization and measures of maturity during the transition from childhood to adolescence have removed large proportions of data (Marek et al., 2019; Dong et al., 2021).

Selection bias could impact analyses of rs-fMRI data collected from such developmental samples if the sample of included children that are able to lay motionless tend to be more mature than the full sample. Diffusion MRI and quantitative susceptibility mapping are also susceptible to motion artifacts (Roalf et al., 2016; He et al., 2015), and as a result, studies using these modalities often exclude participants with gross motion. If quality control procedures in studies using these imaging methods result in a reduced sample in which a variable’s distribution differs from the original sample, and there is evidence that this variable is related to the outcome of interest, then we recommend adjusting means using DRTMLE.

## Supporting information

Web Supplement

## Acknowledgments

The data analyzed in this study were provided in part by grants awarded to SHM from Autism Speaks, the National Institute of Mental Health (R01 MH078160, R01 MH085328), and the National Institute of Neurological Disorders and Stroke (R01 NS048527, R01 NS096207-05); to Richard A. E. Edden from the NIMH (R01 MH106564-03); to Keri Rosch from the NIMH (K23 MH101322-05); to Karen Seymour from the NIMH (K23 MH107734-05); to Bradley Schlagger from the Eunice Kennedy Shriver National Institute of Child Health & Human Development (U54 HD079123). The MRI equipment and resources used to collect the data were provided in part by grants awarded to Peter van Zijl from the NIH (1S10OD021648) and the National Institute of Biomedical Imaging and Bioengineering (P41 EB015909). Analysis, interpretation, and writing of the report were supported by a grant from the NIMH (K01 MH109766 to MBN).

## 6. Citation diversity statement

Recent work in neuroscience (Dworkin et al., 2020) and other fields has identified a citation bias negatively impacting women and other historically underrepresented scholars (Mitchell et al., 2013; Dion et al., 2018; Caplar et al., 2017; Maliniak et al., 2013; Bertolero et al., 2020; Wang et al., 2021; Chatterjee and Werner, 2021; Fulvio et al., 2021). Before writing this manuscript, we set an intention of selecting references that reflect the diversity of the neuroscience and statistics fields in the form of contribution, gender, race, and ethnicity. First, we obtained predicted gender of the first and last authors of each reference using databases that store the probability of a name being carried by a woman or a man (Dworkin et al., 2020; Zhou et al., 2020), with possible combinations including male/male, male/female, female/male, and female/female. Our references contain 14.3% woman(first)/woman(last), 10.9% man/woman, 21.9% woman/man, and 52.9% man/man. man/man. Relative to the expected proportions in the field of neuroscience, we over- or under-cited these categories by the following ratios: 7.6%, 1.5%, -3.6%, and -5.5%, respectively. Second, we obtained the predicted racial/ethnic category of the first and last author of each reference by databases that store the probability of a first and last name being carried by an author of color (Ambekar et al., 2009; Sood and Laohaprapanon, 2018). Our references contain 7.1% author of color (first)/author of color(last), 12.3% white author/author of color, 17.4% author of color/white author, and 63.2% white author/white author. Self citations for the first and last author of the current paper, as well as references for this diversity statement were excluded from these proportion calculations. These methods are limited by the databases and assumptions about gender identity and race they use for prediction, but we look forward to future work that could help us to better understand how to support equitable practices in science.

## CRediT authorship contribution statement

**Mary Beth Nebel**: Conceptualization, Methodology, Software, Validation, Formal Analysis, Data Curation, Writing-Original Draft, Writing-Reviewing & Editing, Visualization, Supervision, Project Administration. **Daniel Lidstone**: Formal Analysis, Writing-Original Draft, Writing-Reviewing & Editing, Visualization. **Liwei Wang**: Software, Formal Analysis, Data Curation, Visualization. **David Benkeser**: Methodology, Software, Writing-Original Draft, Writing-Reviewing & Editing. **Stewart H. Mostofsky**: Conceptualization, Resources, Writing-Review & Editing, Supervision, Project Administration, Funding Acquisition. **Benjamin B. Risk**: Conceptualization, Methodology, Software, Formal Analysis, Writing-Original Draft, Writing-Review & Editing, Visualization, Supervision, Project Administration.

## References

Allen, E. A., Erhardt, E. B., Damaraju, E., Gruner, W., Segall, J. M., Silva, R. F., Havlicek, M., Rachakonda, S., Fries, J., Kalyanam, R., Michael, A. M., Caprihan, A., Turner, J. A., Eichele, T., Adelsheim, S., Bryan, A. D., Bustillo, J., Clark, V. P., Ewing, S. W., Filbey, F., Ford, C. C., Hutchison, K., Jung, R. E., Kiehl, K. A., Kodituwakku, P., Komesu, Y. M., Mayer, A. R., Pearlson, G. D., Phillips, J. P., Sadek, J. R., Stevens, M., Teuscher, U., Thoma, R. J., and Calhoun, V. D. (2011). A baseline for the multivariate comparison of resting-state networks. Frontiers in Systems Neuroscience, 5.

Ambekar, A., Ward, C., Mohammed, J., Male, S., and Skiena, S. (2009). Name-ethnicity classification from open sources. In Proceedings of the 15th ACM SIGKDD international conference on Knowledge Discovery and Data Mining, pages 49–58.

American Psychiatric Association (2013). Diagnostic and statistical manual of mental disorders, 5th ed. (DSM-5®). American Psychiatric Association, 5th edition edition.

Azur, M. J., Stuart, E. A., Frangakis, C., and Leaf, P. J. (2011). Multiple imputation by chained equations: what is it and how does it work? International Journal of Methods in Psychiatric Research, 20(1):40–49.

Bang, H. and Robins, J. M. (2005). Doubly robust estimation in missing data and causal inference models. Biometrics, 61(4):962–973.

Barber, R. F. and Candès, E. J. (2015). Controlling the false discovery rate via knockoffs. Annals of Statistics, 43(5):2055–2085.

Benjamini, Y. and Hochberg, Y. (1995). Controlling the false discovery rate: a practical and powerful approach to multiple testing. Journal of the Royal Statistical Society, Series B (Methodological), 57(1):289–300.

Benkeser, D., Carone, M., van der Laan, M. J., and Gilbert, P. B. (2017). Doubly robust nonparametric inference on the average treatment effect. Biometrika, 104(4):863–880.

Benkeser, D., Petersen, M., and van der Laan, M. J. (2019). Improved Small-Sample Estimation of Nonlinear Cross-Validated Prediction Metrics. Journal of the American Statistical Association, 115(532):1917–1932.

Bertolero, M. A., Dworkin, J. D., David, S. U., Lloreda, C. L., Srivastava, P., Stiso, J., Zhou, D., Dzirasa, K., Fair, D. A., Kaczkurkin, A. N., Marlin, B. J., Shohamy, D., Uddin, L. Q., Zurn, P., and Bassett, D. S. (2020). Racial and ethnic imbalance in neuroscience reference lists and intersections with gender. bioRxiv.

Bijsterbosch, J., Harrison, S. J., Jbabdi, S., Woolrich, M., Beckmann, C., Smith, S., and Duff, E. P. (2020). Challenges and future directions for representations of functional brain organization. Nature Neuroscience, 23(12):1484–1495.

Biswal, B., Zerrin Yetkin, F., Haughton, V. M., and Hyde, J. S. (1995). Functional connectivity in the motor cortex of resting human brain using echo-planar MRI. Magnetic Resonance in Medicine, 34(4):537–541.

Blakemore, S. J., Frith, C. D., and Wolpert, D. M. (2001). The cerebellum is involved in predicting the sensory consequences of action. Neuroreport, 12(9):1879–1884.

Bradley, V. and Nichols, T. E. (2022). Addressing selection bias in the UK Biobank neurological imaging cohort. medRxiv, (2017):2022.01.13.22269266.

Bridgeford, E. W., Powell, M., Kiar, G., Lawrence, R., Caffo, B., Milham, M., and Vogelstein, J. T. (2021). Batch Effects are Causal Effects: Applications in Human Connectomics. bioRxiv, page 2021.09.03.458920.

Buckner, R. L., Krienen, F. M., Castellanos, A., Diaz, J. C., and Thomas Yeo, B. T. (2011). The organization of the human cerebellum estimated by intrinsic functional connectivity. Journal of Neurophysiology, 106(5):2322–2345.

Calhoun, V. D., Adali, T., Pearlson, G. D., and Pekar, J. J. (2001). A method for making group inferences from functional MRI data using independent component analysis. Human Brain Mapping, 14(3):140–151.

Calhoun, V. D., Wager, T. D., Krishnan, A., Rosch, K. S., Seymour, K. E., Nebel, M. B., Mostofsky, S. H., Nyalakanai, P., and Kiehl, K. (2017). The impact of T1 versus EPI spatial normalization templates for fMRI data analyses. Human Brain Mapping, 38(11):5331– 5342.

Caplar, N., Tacchella, S., and Birrer, S. (2017). Quantitative evaluation of gender bias in astronomical publications from citation counts. Nature Astronomy, 1(6):0141.

Casey, B. J., Cannonier, T., Conley, M. I., Cohen, A. O., Barch, D. M., Heitzeg, M. M., Soules, M. E., Teslovich, T., Dellarco, D. V., Garavan, H., Orr, C. A., Wager, T. D., Banich, M. T., Speer, N. K., Sutherland, M. T., Riedel, M. C., Dick, A. S., Bjork, J. M., Thomas, K. M., Chaarani, B., Mejia, M. H., Hagler, D. J., Daniela Cornejo, M., Sicat, C. S., Harms, M. P., Dosenbach, N. U. F., Rosenberg, M., Earl, E., Bartsch, H., Watts, R., Polimeni, J. R., Kuperman, J. M., Fair, D. A., and Dale, A. M. (2018). The Adolescent Brain Cognitive Development (ABCD) study: Imaging acquisition across 21 sites. Developmental Cognitive Neuroscience, 32:43–54.

Chatterjee, P. and Werner, R. M. (2021). Gender disparity in citations in high-impact journal articles. JAMA Netw Open, 4(7):e2114509.

Chen, T. and Guestrin, C. (2016). XGBoost: A Scalable Tree Boosting System. Proceedings of the ACM SIGKDD International Conference on Knowledge Discovery and Data Mining, 13-17-August-2016:785–794.

Ciric, R., Wolf, D. H., Power, J. D., Roalf, D. R., Baum, G. L., Ruparel, K., Shinohara, R. T., Elliott, M. A., Eickhoff, S. B., Davatzikos, C., Gur, R. C., Gur, R. E., Bassett, D. S., and Satterthwaite, T. D. (2017). Benchmarking of participant-level confound regression strategies for the control of motion artifact in studies of functional connectivity. NeuroImage, 154:174–187.

Conners, C. K. (1999). Conners Rating Scales-Revised. In The use of psychological testing for treatment planning and outcomes assessment, 2nd ed, pages 467–495. Lawrence Erlbaum Associates Publishers, Mahwah, NJ, US.

Conners, C. K. (2008). Conners 3 . Multi-Health Systems, Inc, Toronto.

Constantino, J. N. and Gruber, C. P. (2012). Social Responsiveness Scale Second Edition (SRS-2): Manual. Western Psychological Services (WPS).

Constantino, J. N. and Todd, R. D. (2003). Autistic traits in the general population: A twin study. Archives of General Psychiatry, 60(5):524–530.

Crasta, J. E., Zhao, Y., Seymour, K. E., Suskauer, S. J., Mostofsky, S. H., and Rosch, K. S. (2021). Developmental trajectory of subtle motor signs in attention-deficit/hyperactivity disorder: A longitudinal study from childhood to adolescence. Child Neuropsychology, 27(3):317–332.

Dajani, D. R. and Uddin, L. Q. (2016). Local brain connectivity across development in autism spectrum disorder: A cross-sectional investigation. Autism Research, 9(1):43–54.

DeBruine, L. (2018). Plot Comparison [blog post]. retrieved from https://debruine.github.io/post/plot-comparison on 31/10/2021.

Deen, B. and Pelphrey, K. (2012). Perspective: Brain scans need a rethink. Nature, 491(7422 SUPPL.):S20–S20.

Denckla, M. B. (1985). Revised Neurological Examination for Subtle Signs. Psychopharmacology Bulletin, 21(4):773–800.

Di Martino, A., Kelly, C., Grzadzinski, R., Zuo, X. N., Mennes, M., Mairena, M. A., Lord, C., Castellanos, F. X., and Milham, M. P. (2011). Aberrant striatal functional connectivity in children with autism. Biological Psychiatry, 69(9):847–856.

Di Martino, A., O’Connor, D., Chen, B., Alaerts, K., Anderson, J., Assaf, M., Balsters, J., Baxter, L., Beggiato, A., Bernaerts, S., Blanken, L., Bookheimer, S., Braden, B., Byrge, L., Castellanos, F., Dapretto, M., Delorme, R., Fair, D., Fishman, I., Fitzgerald, J., Gallagher, L., Keehn, R., Kennedy, D., Lainhart, J., Luna, B., Mostofsky, S., Müller, R.-A., Nebel, M. B., Nigg, J., O’Hearn, K., Solomon, M., Toro, R., Vaidya, C., Wenderoth, N., White, T., Craddock, R., Lord, C., Leventhal, B., and Milham, M. P. (2017). Enhancing studies of the connectome in autism using the autism brain imaging data exchange II. Scientific Data, 4(1):1–15.

Di Martino, A., Yan, C.-G., Li, Q., Denio, E., Castellanos, F. X., Alaerts, K., Anderson, J. S., Assaf, M., Bookheimer, S., Dapretto, M., Deen, B., Delmonte, S., Dinstein, I., ErtlWagner, B., Fair, D., Gallagher, L., Kennedy, D., Keown, C. L., Keysers, C., Lainhart, J. E., Lord, C., Luna, B., Menon, V., Minshew, N. J., Monk, C., Mueller, S., Müller, R.-A., Nebel, M. B., Nigg, J. T., O’Hearn, K., Pelphrey, K. A., Peltier, S. J., Rudie, J. D., Sunaert, S., Thioux, M., Tyszka, J. M., Uddin, L. Q., Verhoeven, J. S., Wenderoth, N., Wiggins, J. L., Mostofsky, S. H., and Milham, M. P. (2014). The autism brain imaging data exchange: Towards a large-scale evaluation of the intrinsic brain architecture in autism. Molecular Psychiatry, 19(6):659–667.

Dion, M. L., Sumner, J. L., and Mitchell, S. M. (2018). Gendered citation patterns across political science and social science methodology fields. Political Analysis, 26(3):312–327.

Dong, H.-M., Margulies, D. S., Zuo, X.-N., and Holmes, A. J. (2021). Shifting gradients of macroscale cortical organization mark the transition from childhood to adolescence. Proceedings of the National Academy of Sciences, 118(28).

Dosenbach, N. U., Koller, J. M., Earl, E. A., Miranda-Dominguez, O., Klein, R. L., Van, A. N., Snyder, A. Z., Nagel, B. J., Nigg, J. T., Nguyen, A. L., Wesevich, V., Greene, D. J., and Fair, D. A. (2017). Real-time motion analytics during brain MRI improve data quality and reduce costs. NeuroImage, 161:80–93.

D’Souza, N. S., Nebel, M. B., Crocetti, D., Robinson, J., Wymbs, N., Mostofsky, S. H., and Venkataraman, A. (2021). Deep sr-DDL: Deep structurally regularized dynamic dictionary learning to integrate multimodal and dynamic functional connectomics data for multidimensional clinical characterizations. NeuroImage, 241:118388.

DuPaul, G. J., Power, T. J., Anastopoulos, A. D., and Reid, R. (1998). ADHD Rating Scale—IV: Checklists, norms, and clinical interpretation. Guilford Press.

Dworkin, J. D., Linn, K. A., Teich, E. G., Zurn, P., Shinohara, R. T., and Bassett, D. S. (2020). The extent and drivers of gender imbalance in neuroscience reference lists. Nature neuroscience, 23(8):918–926.

Erhardt, E. B., Rachakonda, S., Bedrick, E. J., Allen, E. A., Adali, T., and Calhoun, V. D. (2011). Comparison of multi-subject ICA methods for analysis of fMRI data. Human Brain Mapping, 32(12):2075–2095.

Fassbender, C., Mukherjee, P., and Schweitzer, J. B. (2017). Reprint of: Minimizing noise in pediatric task-based functional MRI; Adolescents with developmental disabilities and typical development. NeuroImage, 154:230–239.

Finn, E. S. and Bandettini, P. A. (2021). Movie-watching outperforms rest for functional connectivity-based prediction of behavior. NeuroImage, 235:117963.

Fortin, J. P., Cullen, N., Sheline, Y. I., Taylor, W. D., Aselcioglu, I., Cook, P. A., Adams, P., Cooper, C., Fava, M., McGrath, P. J., McInnis, M., Phillips, M. L., Trivedi, M. H., Weissman, M. M., and Shinohara, R. T. (2018). Harmonization of cortical thickness measurements across scanners and sites. NeuroImage, 167:104–120.

Friedman, J., Hastie, T., and Tibshirani, R. (2010). Regularization Paths for Generalized Linear Models via Coordinate Descent. Journal of Statistical Software, 33(1):1–22.

Fry, A., Littlejohns, T. J., Sudlow, C., Doherty, N., Adamska, L., Sprosen, T., Collins, R., and Allen, N. E. (2017). Comparison of Sociodemographic and Health-Related Characteristics of UK Biobank Participants With Those of the General Population. American Journal of Epidemiology, 186(9):1026–1034.

Fulvio, J. M., Akinnola, I., and Postle, B. R. (2021). Gender (im)balance in citation practices in cognitive neuroscience. J Cogn Neurosci, 33(1):3–7.

Gabrielsen, T. P., Anderson, J. S., Stephenson, K. G., Beck, J., King, J. B., Kellems, R., Top, D. N., Russell, N. C., Anderberg, E., Lundwall, R. A., Hansen, B., and South, M. (2018). Functional MRI connectivity of children with autism and low verbal and cognitive performance. Molecular Autism, 9(1):1–14.

Greene, D. J., Koller, J. M., Hampton, J. M., Wesevich, V., Van, A. N., Nguyen, A. L., Hoyt, C. R., McIntyre, L., Earl, E. A., Klein, R. L., Shimony, J. S., Petersen, S. E., Schlaggar, B. L., Fair, D. A., and Dosenbach, N. U. (2018). Behavioral interventions for reducing head motion during MRI scans in children. NeuroImage, 171:234–245.

Greenland, S., Robins, J. M., and Pearl, J. (1999). Confounding and collapsibility in causal inference. Statistical Science, 14(1):29–46.

Gu, Z., Gu, L., Eils, R., Schlesner, M., and Brors, B. (2014). circlize implements and enhances circular visualization in R. Bioinformatics, 30(19):2811–2812.

Hastie, T. (2020). gam. Generalized additive models. R package version 1.20 https://CRAN.R-project.org/package-gam.

He, N., Ling, H., Ding, B., Huang, J., Zhang, Y., Zhang, Z., Liu, C., Chen, K., and Yan, F. (2015). Region-specific disturbed iron distribution in early idiopathic Parkinson’s disease measured by quantitative susceptibility mapping. Human Brain Mapping, 36(11):4407– 4420.

Hernan, M. A. and Robins, J. M. (2020). Causal Inference: What If. Chapman Hall/CRC, Boca Raton.

Hollingshead, A. B. (1975). Four factor index of social status. Yale Journal of Sociology, 8.

Horien, C., Fontenelle, S., Joseph, K., Powell, N., Nutor, C., Fortes, D., Butler, M., Powell, K., Macris, D., Lee, K., Greene, A. S., McPartland, J. C., Volkmar, F. R., Scheinost, D., Chawarska, K., and Constable, R. T. (2020). Low-motion fMRI data can be obtained in pediatric participants undergoing a 60-minute scan protocol. Scientific Reports 2020 *10*:1, 10(1):1–10.

Hus, V., Gotham, K., and Lord, C. (2014). Standardizing ADOS domain scores: Separating severity of social affect and restricted and repetitive behaviors. Journal of Autism and Developmental Disorders, 44(10):2400–2412.

Jenkinson, M., Bannister, P., Brady, M., and Smith, S. (2002). Improved Optimization for the Robust and Accurate Linear Registration and Motion Correction of Brain Images. NeuroImage, 17:825–841.

Johnstone, T., Ores Walsh, K. S., Greischar, L. L., Alexander, A. L., Fox, A. S., Davidson, R. J., and Oakes, T. R. (2006). Motion correction and the use of motion covariates in multiple-subject fMRI analysis. Human Brain Mapping, 27(10):779–788.

Jones, T. B., Bandettini, P. A., Kenworthy, L., Case, L. K., Milleville, S. C., Martin, A., and Birn, R. M. (2010). Sources of group differences in functional connectivity: An investigation applied to autism spectrum disorder. NeuroImage, 49(1):401–414.

Kaufman, J., Birmaher, B., Axelson, D., Perepletchikova, F., Brent, D., and Ryan, N. (2013). Schedule for affective disorders and schizophrenia for school-age children-present and lifetime version (K-SADS-PL 2013, DSM-5). Pittsburgh, PA: Western Psychiatric Institute and Clinic and Yale University.

Kennedy, D. P., Redcay, E., and Courchesne, E. (2006). Failing to deactivate: Resting functional abnormalities in autism. Proceedings of the National Academy of Sciences of the United States of America, 103(21):8275–8280.

Keown, C. L., Shih, P., Nair, A., Peterson, N., Mulvey, M. E., and Müller, R. A. (2013). Local functional overconnectivity in posterior brain regions is associated with symptom severity in autism spectrum disorders. Cell Reports, 5(3):567–572.

Kong, X. Z., Zhen, Z., Li, X., Lu, H. H., Wang, R., Liu, L., He, Y., Zang, Y., and Liu, J. (2014). Individual differences in impulsivity predict head motion during magnetic resonance imaging. PLoS ONE, 9(8):e104989.

Lake, E. M., Finn, E. S., Noble, S. M., Vanderwal, T., Shen, X., Rosenberg, M. D., Spann, M. N., Chun, M. M., Scheinost, D., and Constable, R. T. (2019). The functional brain organization of an individual allows prediction of measures of social abilities transdiagnostically in autism and attention-deficit/hyperactivity disorder. Biological Psychiatry, 86(4):315–326.

Lash, T. L., VanderWeele, T. J., Haneuse, S., and Rothman, K. J. (2021). Modern Epidemiology, Fourth Edition. Wolters Kluwer.

Lewinn, K. Z., Sheridan, M. A., Keyes, K. M., Hamilton, A., and McLaughlin, K. A. (2017). Sample composition alters associations between age and brain structure. Nature Communications, 8(1):1–14.

Lidstone, D. E., Rochowiak, R., Mostofsky, S. H., and Nebel, M. B. (2021). A Data Driven Approach Reveals That Anomalous Motor System Connectivity is Associated With the Severity of Core Autism Symptoms. Autism Research, Epub ahead of print:1–18.

Little, R. J. and Rubin, D. B. (2019). Statistical analysis with missing data, volume 793. John Wiley & Sons.

Lombardo, M. V., Eyler, L., Moore, A., Datko, M., Barnes, C. C., Cha, D., Courchesne, E., and Pierce, K. (2019). Default mode-visual network hypoconnectivity in an autism subtype with pronounced social visual engagement difficulties. eLife, 8:e47427.

Lord, C. and Jones, R. M. (2012). Annual research review: Re-thinking the classification of autism spectrum disorders. Journal of Child Psychology and Psychiatry and Allied Disciplines, 53(5):490–509.

Lord, C., Risi, S., Lambrecht, L., Cook, E. H., Leventhal, B. L., Dilavore, P. C., Pickles, A., and Rutter, M. (2000). The Autism Diagnostic Observation Schedule-Generic: A standard measure of social and communication deficits associated with the spectrum of autism. Journal of Autism and Developmental Disorders, 30(3):205–223.

Maenner, M. J., Shaw, K. A., Bakian, A. V., Bilder, D. A., Durkin, M. S., Esler, A., Furnier, S. M., Hallas, L., Hall-Lande, J., Hudson, A., Hughes, M. M., Patrick, M., Pierce, K., Poynter, J. N., Salinas, A., Shenouda, J., Vehorn, A., Warren, Z., Constantino, J. N., DiRienzo, M., Fitzgerald, R. T., Grzybowski, A., Spivey, M. H., Pettygrove, S., Zahorodny, W., Ali, A., Andrews, J. G., Baroud, T., Gutierrez, J., Hewitt, A., Lee, L.-C., Lopez, M., Mancilla, K. C., McArthur, D., Schwenk, Y. D., Washington, A., Williams, S., and Cogswell, M. E. (2021). Prevalence and Characteristics of Autism Spectrum Disorder Among Children Aged 8 Years — Autism and Developmental Disabilities Monitoring Network, 11 Sites, United States, 2018. MMWR. Surveillance Summaries, 70(11):1–16.

Mahadevan, A. S., Tooley, U. A., Bertolero, M. A., Mackey, A. P., and Bassett, D. S. (2021). Evaluating the sensitivity of functional connectivity measures to motion artifact in resting-state fMRI data. NeuroImage, 241:118408.

Maliniak, D., Powers, R., and Walter, B. F. (2013). The gender citation gap in international relations. International Organization, 67(4):889–922.

Marek, S., Tervo-Clemmens, B., Calabro, F. J., Montez, D. F., Kay, B. P., Hatoum, A. S., Donohue, M. R., Foran, W., Miller, R. L., Hendrickson, T. J., Malone, S. M., Kandala, S., Feczko, E., Miranda-Dominguez, O., Graham, A. M., Earl, E. A., Perrone, A. J., Cordova, M., Doyle, O., Moore, L. A., Conan, G. M., Uriarte, J., Snider, K., Lynch, B. J., Wilgenbusch, J. C., Pengo, T., Tam, A., Chen, J., Newbold, D. J., Zheng, A., Seider, N. A., Van, A. N., Metoki, A., Chauvin, R. J., Laumann, T. O., Greene, D. J., Petersen, S. E., Garavan, H., Thompson, W. K., Nichols, T. E., Yeo, B. T. T., Barch, D. M., Luna, B., Fair, D. A., and Dosenbach, N. U. F. (2022). Reproducible brain-wide association studies require thousands of individuals. Nature 2022, pages 1–7.

Marek, S., Tervo-Clemmens, B., Nielsen, A. N., Wheelock, M. D., Miller, R. L., Laumann, T. O., Earl, E., Foran, W. W., Cordova, M., Doyle, O., Perrone, A., Miranda-Dominguez, O., Feczko, E., Sturgeon, D., Graham, A., Hermosillo, R., Snider, K., Galassi, A., Nagel, B. J., Ewing, S. W., Eggebrecht, A. T., Garavan, H., Dale, A. M., Greene, D. J., Barch, D. M., Fair, D. A., Luna, B., and Dosenbach, N. U. (2019). Identifying reproducible individual differences in childhood functional brain networks: An ABCD study. Developmental Cognitive Neuroscience, 40:100706.

Mayes, S. D. and Calhoun, S. L. (2008). WISC-IV and WIAT-II profiles in children with high-functioning autism. Journal of Autism and Developmental Disorders, 38(3):428–439.

Mejia, A. F., Nebel, M. B., Barber, A., Choe, A., Pekar, J., Caffo, B., and Lindquist, M. A. (2018). Improved estimation of subject-level functional connectivity using full and partial correlation with empirical Bayes shrinkage. NeuroImage, 172:478–491.

Mejia, A. F., Nebel, M. B., Eloyan, A., Caffo, B., and Lindquist, M. A. (2017). PCA leverage: outlier detection for high-dimensional functional magnetic resonance imaging data. Biostatistics, 18(3):521–536.

Meyer, D., Dimitriadou, E., Hornik, K., Weingessel, A., and Leisch, F. (2021). e1071: Misc functions of the Department of Statistics, Probability Theory Group (Formerly: E1071), TU Wien. *R package version 1.7-9.* https://CRAN-R-project.org/package-e1071.

Milborrow, S. (2011). earth: Multivariate Adaptive Regression Splines. Derived from mda:mars by T. Hastie and R. Tibshirani. R package. http://CRAN.R-project.org/package=earth.

Mitchell, S. M., Lange, S., and Brus, H. (2013). Gendered citation patterns in international relations journals. International Studies Perspectives, 14(4):485–492.

Moons, K. G., Donders, R. A., Stijnen, T., and Harrell, F. E. (2006). Using the outcome for imputation of missing predictor values was preferred. Journal of Clinical Epidemiology, 59(10):1092–1101.

Moore, K. L. and van der Laan, M. J. (2009). Covariate adjustment in randomized trials with binary outcomes: Targeted maximum likelihood estimation. Statistics in Medicine, 28(1):39–64.

Mostofsky, S. H., Newschaffer, C. J., and Denckla, M. B. (2003). Overflow movements predict impaired response inhibition in children with ADHD. Perceptual and Motor Skills, 97(3 II):1315–1331.

Mowinckel, A. (2018). Circular Plots in R and Adding Images [blog post]. retrieved from https://drmowinckels.io/blog/2018-05-25-circluar-plots-in-r-and-adding-images on 31/10/2021.

Muschelli, J., Nebel, M. B., Caffo, B., Barber, A., Pekar, J., and Mostofsky, S. H. (2014). Reduction of motion-related artifacts in resting state fMRI using aCompCor. NeuroImage, 96:22–35.

Nielsen, A. N., Greene, D. J., Gratton, C., Dosenbach, N. U., Petersen, S. E., and Schlaggar, B. L. (2019). Evaluating the prediction of brain maturity from functional connectivity after motion artifact denoising. Cerebral Cortex, 29(6):2455–2469.

Nordahl, C. W., Mello, M., Shen, A. M., Shen, M. D., Vismara, L. A., Li, D., Harrington, K., Tanase, C., Goodlin-Jones, B., Rogers, S., Abbeduto, L., and Amaral, D. G. (2016). Methods for acquiring MRI data in children with autism spectrum disorder and intellectual impairment without the use of sedation. Journal of Neurodevelopmental Disorders, 8(1):1– 10.

Padmanabhan, A., Lynch, C. J., Schaer, M., and Menon, V. (2017). The Default Mode Network in Autism. Biological Psychiatry: Cognitive Neuroscience and Neuroimaging, 2(6):476–486.

Parkes, L., Fulcher, B., Yücel, M., and Fornito, A. (2018). An evaluation of the efficacy, reliability, and sensitivity of motion correction strategies for resting-state functional MRI. NeuroImage, 171:415–436.

Petersen, M. L., Porter, K. E., Gruber, S., Wang, Y., and van der Laan, M. J. (2010). Diagnosing and responding to violations in the positivity assumption:. Statistical Methods in Medical Research, 21(1):31–54.

Polley, E., LeDell, E., Kennedy, C., and van der Laan, M. (2019). SuperLearner: Super Learner Prediction. R package v. 2.0-26.

Power, J. D. (2017). A simple but useful way to assess fMRI scan qualities. NeuroImage, 154:150–158.

Power, J. D., Barnes, K. A., Snyder, A. Z., Schlaggar, B. L., and Petersen, S. E. (2012). Spurious but systematic correlations in functional connectivity MRI networks arise from subject motion. NeuroImage, 59(3):2142–2154.

Power, J. D., Lynch, C. J., Adeyemo, B., and Petersen, S. E. (2020). A critical, event-related appraisal of denoising in resting-state fMRI studies. Cerebral Cortex, 30(10):5544–5559.

Power, J. D., Mitra, A., Laumann, T. O., Snyder, A. Z., Schlaggar, B. L., and Petersen, S. E. (2014). Methods to detect, characterize, and remove motion artifact in resting state fMRI. NeuroImage, 84:320–341.

Pruim, R. H. R., Mennes, M., Buitelaar, J. K., and Beckmann, C. F. (2015). Evaluation of ICA-AROMA and alternative strategies for motion artifact removal in resting state fMRI. NeuroImage, 112:278–287.

Pua, E. P. K., Barton, S., Williams, K., Craig, J. M., and Seal, M. L. (2020). Individualised MRI training for paediatric neuroimaging: A child-focused approach. Developmental Cognitive Neuroscience, 41:100750.

Redcay, E., Moran, J. M., Mavros, P. L., Tager-Flusberg, H., Gabrieli, J. D., and Whitfield-Gabrieli, S. (2013). Intrinsic functional network organization in high-functioning adolescents with autism spectrum disorder. Frontiers in Human Neuroscience, 7:573.

Reich, W. (2000). Diagnostic Interview for Children and Adolescents (DICA). Journal of the American Academy of Child and Adolescent Psychiatry, 39(1):59–66.

Reiter, M. A., Mash, L. E., Linke, A. C., Fong, C. H., Fishman, I., and Müller, R. A. (2019). Distinct Patterns of Atypical Functional Connectivity in Lower-Functioning Autism. Biological Psychiatry: Cognitive Neuroscience and Neuroimaging, 4(3):251–259.

Roalf, D. R., Quarmley, M., Elliott, M. A., Satterthwaite, T. D., Vandekar, S. N., Ruparel, K., Gennatas, E. D., Calkins, M. E., Moore, T. M., Hopson, R., Prabhakaran, K., Jackson, C. T., Verma, R., Hakonarson, H., Gur, R. C., and Gur, R. E. (2016). The impact of quality assurance assessment on diffusion tensor imaging outcomes in a large-scale population-based cohort. NeuroImage, 125:903–919.

Robins, J. (1986). A new approach to causal inference in mortality studies with a sustained exposure period-application to control of the healthy worker survivor effect. Mathematical Modelling, 7(9-12):1393–1512.

Robins, J. M., Rotnitzky, A., and Zhao, L. P. (2012). Estimation of Regression Coefficients When Some Regressors are not Always Observed. https://doi.org/10.1080/01621459.1994.10476818, 89(427):846–866.

Rudie, J. D., Brown, J. A., Beck-Pancer, D., Hernandez, L. M., Dennis, E. L., Thompson, P. M., Bookheimer, S. Y., and Dapretto, M. (2013). Altered functional and structural brain network organization in autism. NeuroImage: Clinical, 2(1):79–94.

Satterthwaite, T. D., Elliott, M. A., Gerraty, R. T., Ruparel, K., Loughead, J., Calkins, M. E., Eickhoff, S. B., Hakonarson, H., Gur, R. C., Gur, R. E., and Wolf, D. H. (2013). An improved framework for confound regression and filtering for control of motion artifact in the preprocessing of resting-state functional connectivity data. NeuroImage, 64:240–256.

Satterthwaite, T. D., Wolf, D. H., Loughead, J., Ruparel, K., Elliott, M. A., Hakonarson, H., Gur, R. C., and Gur, R. E. (2012). Impact of in-scanner head motion on multiple measures of functional connectivity: Relevance for studies of neurodevelopment in youth. NeuroImage, 60(1):623–632.

Schuler, M. S. and Rose, S. (2017). Targeted maximum likelihood estimation for causal inference in observational studies. American Journal of Epidemiology, 185(1):65–73.

Simhal, A. K., Filho, J. O., Segura, P., Cloud, J., Petkova, E., Gallagher, R., Castellanos, F. X., Colcombe, S., Milham, M. P., and Di Martino, A. (2021). Predicting multiscan MRI outcomes in children with neurodevelopmental conditions following MRI simulator training. Developmental Cognitive Neuroscience, 52:101009.

Simonoff, E., Pickles, A., Charman, T., Chandler, S., Loucas, T., and Baird, G. (2008). Psychiatric disorders in children with autism spectrum disorders: Prevalence, comorbidity, and associated factors in a population-derived sample. Journal of the American Academy of Child and Adolescent Psychiatry, 47(8):921–929.

Snowden, J. M., Rose, S., and Mortimer, K. M. (2011). Implementation of G-Computation on a Simulated Data Set: Demonstration of a Causal Inference Technique. American Journal of Epidemiology, 173(7):731–738.

Sood, G. and Laohaprapanon, S. (2018). Predicting race and ethnicity from the sequence of characters in a name. arXiv preprint *arXiv:1805.02109*.

Stoodley, C. J., D’Mello, A. M., Ellegood, J., Jakkamsetti, V., Liu, P., Nebel, M. B., Gibson, J. M., Kelly, E., Meng, F., Cano, C. A., Pascual, J. M., Mostofsky, S. H., Lerch, J. P., and Tsai, P. T. (2017). Altered cerebellar connectivity in autism and cerebellar-mediated rescue of autism-related behaviors in mice. Nature Neuroscience, 20(12):1744–1751.

Stuart, E. A. (2010). Matching methods for causal inference: A review and a look forward. Statistical science : a review journal of the Institute of Mathematical Statistics, 25(1):1–21.

Stunden, C., Stratton, K., Zakani, S., and Jacob, J. M. (2021). Comparing a Virtual Reality-Based Simulation App (VR-MRI) With a Standard Preparatory Manual and Child Life Program for Improving Success and Reducing Anxiety During Pediatric Medical Imaging: Randomized Clinical Trial. Journal of medical Internet research, 23(9).

Supekar, K., Uddin, L. Q., Khouzam, A., Phillips, J., Gaillard, W. D., Kenworthy, L. E., Yerys, B. E., Vaidya, C. J., and Menon, V. (2013). Brain Hyperconnectivity in Children with Autism and its Links to Social Deficits. Cell Reports, 5(3):738–747.

Tyszka, J. M., Kennedy, D. P., Paul, L. K., and Adolphs, R. (2014). Largely typical patterns of resting-state functional connectivity in high-functioning adults with autism. Cerebral Cortex, 24(7):1894–1905.

Uddin, L. Q., Supekar, K., Lynch, C. J., Khouzam, A., Phillips, J., Feinstein, C., Ryali, S., and Menon, V. (2013). Salience network-based classification and prediction of symptom severity in children with autism. JAMA Psychiatry, 70(8):869–879.

Uddin, L. Q., Yeo, B. T., and Spreng, R. N. (2019). Towards a Universal Taxonomy of Macro-scale Functional Human Brain Networks. Brain Topography, 32(6):926–942.

van der Laan, M. J., Polley, E. C., and Hubbard, A. E. (2007). Super learner. Statistical Applications in Genetics and Molecular Biology, 6(1).

van der Laan, M. J. and Robins, J. M. (2003). Unified Methods for Censored Longitudinal Data and Causality. Springer Science & Business Media.

van der Laan, M. J. and Rose, S. (2011). Targeted Learning: Causal Inference for Observational and Experimental Data. Springer, New York, NY.

van der Laan, M. J. and Rubin, D. (2006). Targeted maximum likelihood learning. International Journal of Biostatistics, 2(1).

van Dijk, K. R., Sabuncu, M. R., and Buckner, R. L. (2012). The influence of head motion on intrinsic functional connectivity MRI. NeuroImage, 59(1):431–438.

Van Overwalle, F., Van de Steen, F., van Dun, K., and Heleven, E. (2020). Connectivity between the cerebrum and cerebellum during social and non-social sequencing using dynamic causal modelling. NeuroImage, 206:116326.

Vander Weele, T. J. (2009). Concerning the consistency assumption in causal inference. Epidemiology, 20(6):880–883.

Vanderwal, T., Eilbott, J., and Castellanos, F. X. (2019). Movies in the magnet: Naturalistic paradigms in developmental functional neuroimaging.

Vasa, R. A., Mostofsky, S. H., and Ewen, J. B. (2016). The Disrupted Connectivity Hypothesis of Autism Spectrum Disorders: Time for the Next Phase in Research. Biological Psychiatry: Cognitive Neuroscience and Neuroimaging, 1(3):245–252.

Venables, W. N. and Ripley, B. D. (2002). Modern Applied Statistics with S. Statistics and Computing. Springer New York, New York, NY, 4th edition.

Wang, X., Dworkin, J., Zhou, D., Stiso, J., Falk, E. B., Bassett, D., Zurn, P., and Lydon-Staley, D. M. (2021). Gendered citation practices in the field of communication. Annals of the International Communication Association, 45(2):134–153.

Wechsler, D. (2003). Wechsler Intelligence Scale for Children-WISC-IV. Psychological Corporation.

Wickham, H. (2016). ggplot2: Elegant Graphics for Data Analysis. Springer-Verlag New York.

Winkler, A. M., Ridgway, G. R., Webster, M. A., Smith, S. M., and Nichols, T. E. (2014). Permutation inference for the general linear model. NeuroImage, 92(100):381–397.

Wood, S. N. (2017). Generalized additive models: an introduction with R. CRC Press.

Wright, M. N. and Ziegler, A. (2017). ranger: A Fast Implementation of Random Forests for High Dimensional Data in C++ and R. Journal of Statistical Software, 77(1):1–17.

Wymbs, N. F., Nebel, M. B., Ewen, J. B., and Mostofsky, S. H. (2021). Altered inferior parietal functional connectivity is correlated with praxis and social skill performance in children with autism spectrum disorder. Cerebral Cortex, 31(5):2639–2652.

Xie, S., Karlsson, H., Dalman, C., Widman, L., Rai, D., Gardner, R. M., Magnusson, C., Sandin, S., Tabb, L. P., Newschaffer, C. J., and Lee, B. K. (2020). The Familial Risk of Autism Spectrum Disorder with and without Intellectual Disability. Autism Research, 13(12):2242–2250.

Yan, C. G., Cheung, B., Kelly, C., Colcombe, S., Craddock, R. C., Di Martino, A., Li, Q., Zuo, X. N., Castellanos, F. X., and Milham, M. P. (2013). A comprehensive assessment of regional variation in the impact of head micromovements on functional connectomics. NeuroImage, 76:183–201.

Zhou, D., Cornblath, E. J., Stiso, J., Teich, E. G., Dworkin, J. D., Blevins, A. S., and Bassett, D. S. (2020). Gender diversity statement and code notebook v1.0 [software]. retrieved from https://github.com/dalejn/cleanBib *on 30/11*/2021.

