## Supplementary material for "Accounting for motion in resting-state fMRI: What part of the spectrum are we characterizing in autism spectrum disorder?": Web Supplement

Table S.1: **Summary of Mann-Whitney U tests comparing included and excluded participants using the lenient motion QC and stratified by primary diagnosis.** Medians for included and excluded participants are reported, as well as effect sizes (r) and FDR-adjusted p values. ASD=autism spectrum disorder. ADOS=Autism Diagnostic Observation Schedule total score. GAI=General Ability Index. SRS=Social Responsiveness Scale. TD=typically developing

| Primary Diagnosis | Variable | Included | Excluded | r | p |
|---|---|---|---|---|---|
| ASD | ADOS | 13.00 | 16.00 | 0.16 | 0.07 |
| ASD | SRS | 91.25 | 100.00 | 0.19 | 0.05 |
| ASD | Motor Overflow | 17.00 | 22.00 | 0.27 | 0.01 |
| ASD | Inattention | 18.00 | 16.00 | 0.01 | 0.75 |
| ASD | Hyperactivity | 11.00 | 12.00 | 0.03 | 0.42 |
| ASD | Age | 10.54 | 9.76 | 0.14 | 0.08 |
| ASD | GAI | 108.00 | 101.00 | 0.15 | 0.07 |
| TD | SRS | 16.00 | 15.00 | <0.01 | 0.54 |
| TD | Motor Overflow | 11.00 | 13.00 | 0.08 | 0.08 |
| TD | Inattention | 2.00 | 2.00 | 0.03 | 0.34 |
| TD | Hyperactivity | 1.00 | 2.00 | 0.11 | 0.07 |
| TD | Age | 10.32 | 9.87 | 0.12 | 0.05 |
| TD | GAI | 116.00 | 112.50 | 0.10 | 0.07 |

*716 N Broadway, Baltimore, MD 21205

Table S.2: **Summary of Mann-Whitney U tests comparing included and excluded participants using the strict motion QC and stratified by primary diagnosis.** Medians for included and excluded participants are reported, as well as effect sizes (r) and FDR-adjusted p values. ASD=autism spectrum disorder. ADOS=Autism Diagnostic Observation Schedule total score. GAI=General Ability Index. SRS=Social Responsiveness Scale. TD=typically developing

| Primary Diagnosis | variable | Included | Excluded | r | p |
|---|---|---|---|---|---|
| ASD | ADOS | 12.50 | 14.00 | 0.17 | 0.08 |
| ASD | SRS | 84.25 | 93.50 | 0.17 | 0.08 |
| ASD | Motor Overflow | 15.00 | 18.00 | 0.15 | 0.10 |
| ASD | Inattention | 15.50 | 18.00 | 0.06 | 0.25 |
| ASD | Hyperactivity | 11.00 | 12.00 | 0.07 | 0.25 |
| ASD | Age | 11.04 | 10.13 | 0.20 | 0.05 |
| ASD | GAI | 107.50 | 107.00 | 0.02 | 0.40 |
| TD | SRS | 14.25 | 17.00 | 0.10 | 0.10 |
| TD | Motor Overflow | 11.00 | 12.00 | 0.06 | 0.20 |
| TD | Inattention | 2.00 | 2.00 | 0.05 | 0.20 |
| TD | Hyperactivity | 1.00 | 2.00 | 0.16 | 0.02 |
| TD | Age | 10.38 | 10.13 | 0.09 | 0.10 |
| TD | GAI | 116.00 | 115.00 | 0.07 | 0.14 |

Table S.3: **Summary of edges for which DRTMLE indicated a group difference at FDR=.20**. Naïve group mean functional connectivity estimates, naïve Z-Statistics for ASD-TD, DRTMLE group mean functional connectivity estimates, DRTMLE Z-Statistics for ASD-TD, naïve FDR-adjusted p-values, and DRTMLE FDR-adjusted p-values are displayed. Mean functional connectivity group estimates further from zero reflect stronger functional connectivity regardless of sign. Positive mean functional connectivity estimate reflect positive partial correlations, or more integrated intrinsic activity between independent components (ICs). Negative mean functional connectivity estimates reflect negative partial correlations or more segregated intrinsic activity between ICs. Edges are ordered by naïve FDR-adjusted p-values. ASD=autism spectrum disorder. TD=typically developing. Attn=dorsal attention. CB=pontomedullary/cerebellar. Control=executive control. Default=default mode. SalVenAtt=salience/ventral attention. SomMot=somatomotor. Vis=visual.

| Edge | Network Pair | naive ASD mean | naive TD mean | naive ASD-TD Z | DRTMLE ASD mean | DRTMLE TD mean | DRTMLE ASD-TD Z | naive FDR-adjusted p | DRTMLE FDR-adjusted p |
|---|---|---|---|---|---|---|---|---|---|
| 2-27 | Vis-Control | -0.038 | -0.024 | -4.088 | -0.040 | -0.024 | -4.67 | 0.007 | 0.000 |
| 14-19 | CB-Attn | 0.068 | 0.078 | -3.351 | 0.068 | 0.078 | -3.18 | 0.062 | 0.045 |
| 13-26 | SomMot-Default | -0.031 | -0.039 | 2.805 | -0.030 | -0.039 | 3.12 | 0.128 | 0.046 |
| 14-21 | CB-Control | 0.044 | 0.053 | -2.862 | 0.041 | 0.052 | -3.36 | 0.128 | 0.045 |
| 19-25 | Attn-Default | -0.085 | -0.094 | 2.867 | -0.086 | -0.094 | 2.45 | 0.128 | 0.168 |
| 19-26 | Attn-Default | 0.037 | 0.028 | 2.997 | 0.037 | 0.027 | 3.29 | 0.128 | 0.045 |
| 4-17 | CB-Default | -0.067 | -0.076 | 2.671 | -0.067 | -0.076 | 3.19 | 0.165 | 0.045 |
| 17-24 | Default-Control | -0.005 | 0.003 | -2.561 | -0.006 | 0.003 | -2.94 | 0.200 | 0.063 |
| 1-14 | SomMot-CB | -0.002 | 0.007 | -2.225 | -0.003 | 0.006 | -2.41 | 0.399 | 0.168 |
| 14-24 | CB-Control | 0.015 | 0.021 | -2.099 | 0.014 | 0.021 | -2.30 | 0.457 | 0.168 |
| 21-30 | Control-SalVenAtt | 0.063 | 0.069 | -2.115 | 0.063 | 0.069 | -2.33 | 0.457 | 0.168 |
| 13-25 | SomMot-Default | -0.097 | -0.102 | 1.869 | -0.094 | -0.102 | 2.66 | 0.470 | 0.109 |
| 15-30 | SomMot-SalVenAtt | 0.113 | 0.117 | -1.830 | 0.112 | 0.117 | -2.29 | 0.470 | 0.168 |
| 19-24 | Attn-Control | -0.160 | -0.165 | 2.037 | -0.159 | -0.165 | 2.25 | 0.470 | 0.171 |
| 4-21 | CB-Control | -0.041 | -0.034 | -1.918 | -0.043 | -0.034 | -2.17 | 0.470 | 0.184 |
| 8-22 | Vis-SalVenAtt | 0.019 | 0.024 | -1.782 | 0.017 | 0.025 | -2.48 | 0.470 | 0.168 |
| 26-30 | Default-SalVenAtt | -0.088 | -0.082 | -1.720 | -0.090 | -0.082 | -2.29 | 0.503 | 0.168 |
| 1-24 | SomMot-Control | -0.027 | -0.032 | 1.695 | -0.023 | -0.033 | 2.74 | 0.510 | 0.094 |
| 1-21 | SomMot-Control | 0.003 | -0.002 | 1.668 | 0.008 | -0.002 | 3.02 | 0.521 | 0.055 |
| 13-22 | SomMot-SalVenAtt | -0.030 | -0.035 | 1.564 | -0.029 | -0.035 | 2.35 | 0.522 | 0.168 |
| 17-27 | Default-Control | 0.034 | 0.028 | 1.557 | 0.036 | 0.028 | 2.17 | 0.522 | 0.184 |
| 13-24 | SomMot-Control | -0.091 | -0.095 | 1.117 | -0.088 | -0.095 | 2.24 | 0.679 | 0.171 |
| 15-21 | SomMot-Control | 0.039 | 0.042 | -1.098 | 0.038 | 0.044 | -2.23 | 0.679 | 0.171 |
| 19-21 | Attn-Control | -0.009 | -0.013 | 1.086 | -0.006 | -0.015 | 2.75 | 0.679 | 0.094 |
| 17-25 | Default-Default | 0.033 | 0.032 | 0.425 | 0.041 | 0.031 | 2.31 | 0.916 | 0.168 |

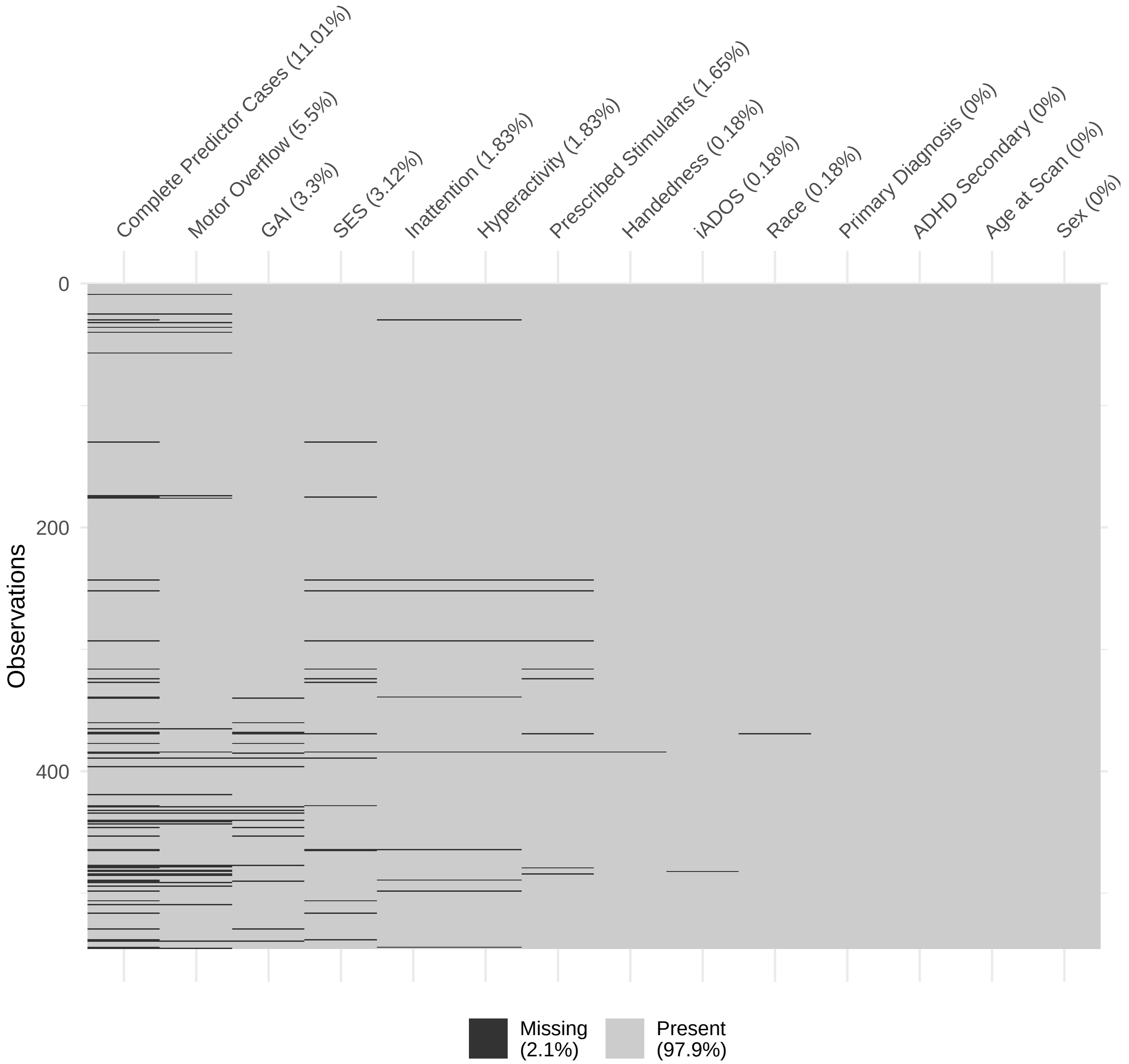


Figure S.1: **Missingness of socio-demographic and behavioral variables in the initial KKI dataset.** There are 545 children in the aggregated KKI dataset. The sample used in DRTMLE is defined by the observations containing data for the variables in the propensity and outcome models together with the socio-demographic variables used in adjusted residuals (i.e., Socioeconomic Status [SES], Sex, and Race). Ordered by the percent missingness, the variables used to define the complete predictor cases are GAI, SES, Motor Overflow, Inattention, Hyperactivity, Prescribed Stimulants, Handedness, Autism Diagnostic Observation Schedule total score including assignment ADOS=0 for typically developing (iADOS), Race, Primary Diagnosis, ADHD Secondary Diagnosis, Age at Scan, and Sex.

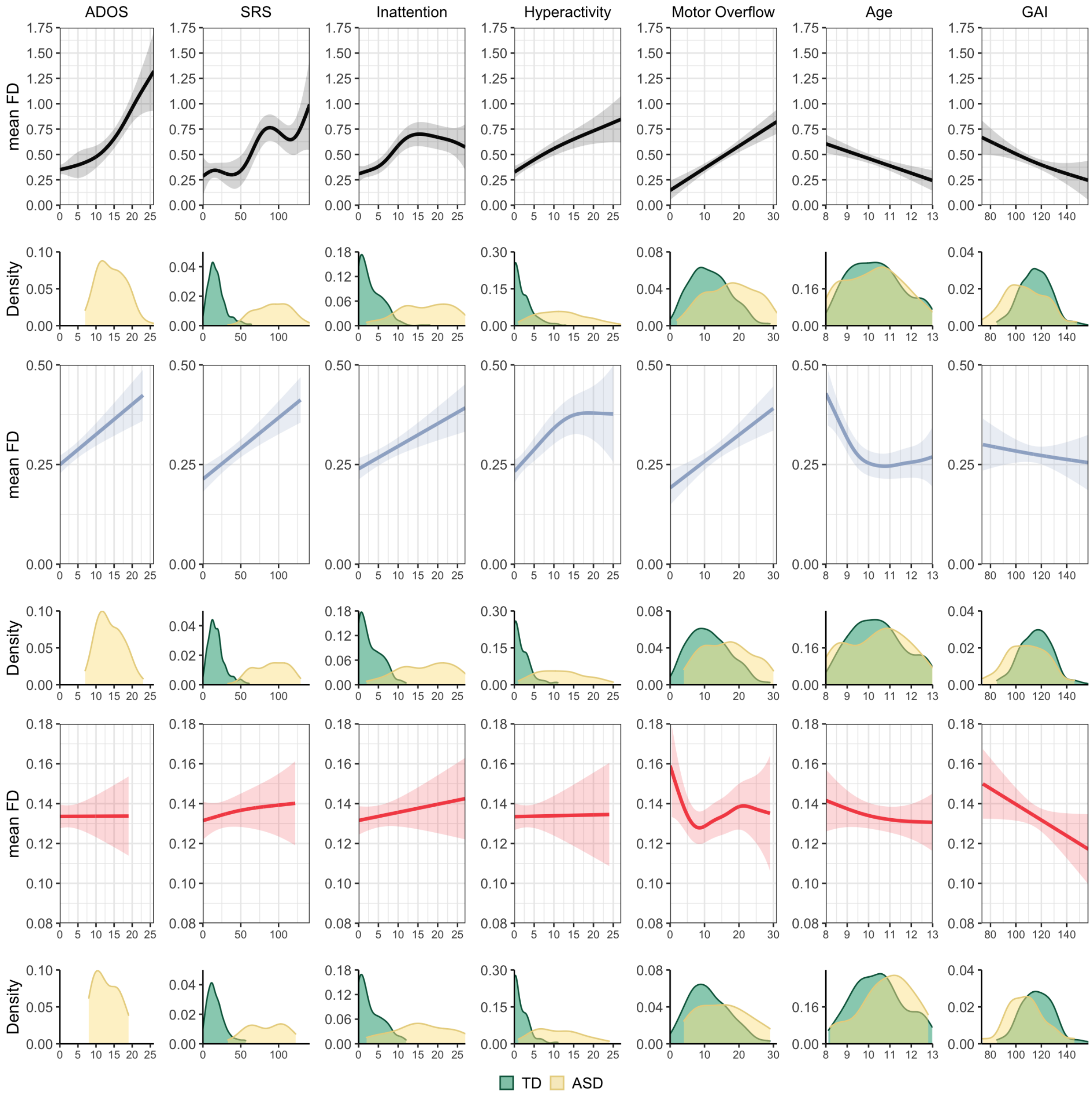


Figure S.2: **Univariate analysis of mean framewise displacement as a function of participant characteristics**. From left to right: Autism Diagnostic Observation Schedule (ADOS) total scores, social responsiveness scale (SRS) scores, inattentive symptoms, hyperactive/impulsive symptoms, total motor overflow, age, and general ability index (GAI) for children with usable and unusable rsfMRI data (top row, black lines), children with usable data under lenient motion QC (slate blue lines), and children with usable data under strict motion QC (red lines). Variable distributions for each diagnosis group are displayed across the bottom panel (TD=typically developing, green; ASD=autism spectrum disorder, yellow).

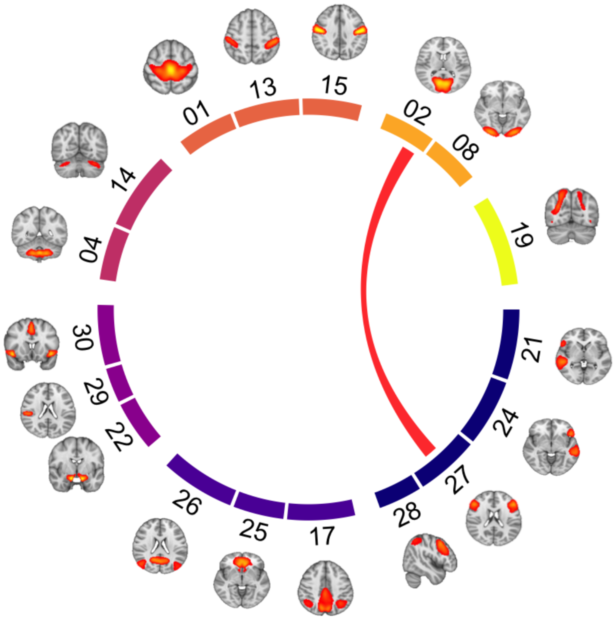


(a) Naïve Z-Statistic

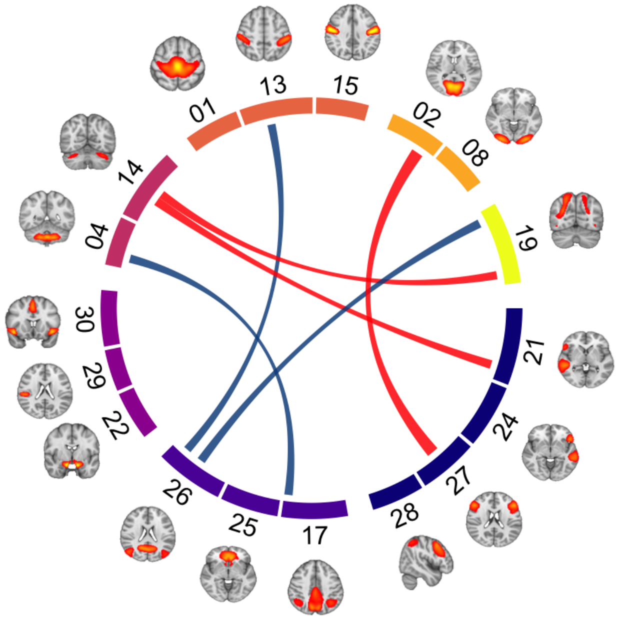


(b) DRTMLE Z-Statistic

Figure S.3: **Comparison of edges showing a group difference using the naïve and DRTMLE approaches at FDR=0.05**. Z-statistics for autism spectrum disorder (ASD) versus typically developing (TD) using A) the naive test and B) the deconfounded group difference using DRTMLE. Connections are thresholded at FDR=0.05. Blue lines indicates ASD>TD (0 in naïve, 3 in DRTMLE). Red lines indicates ASD<TD (1 in naïve, 3 in DRTMLE). Brain regions contributing to each independent component are illustrated and components are grouped by functional assignment. Navy nodes: control. Blue violet: default mode. Purple: salience/ventral attention. Magenta: pontomedullary/cerebellar. Coral: somatomotor. Orange: visual. Yellow: dorsal attention.

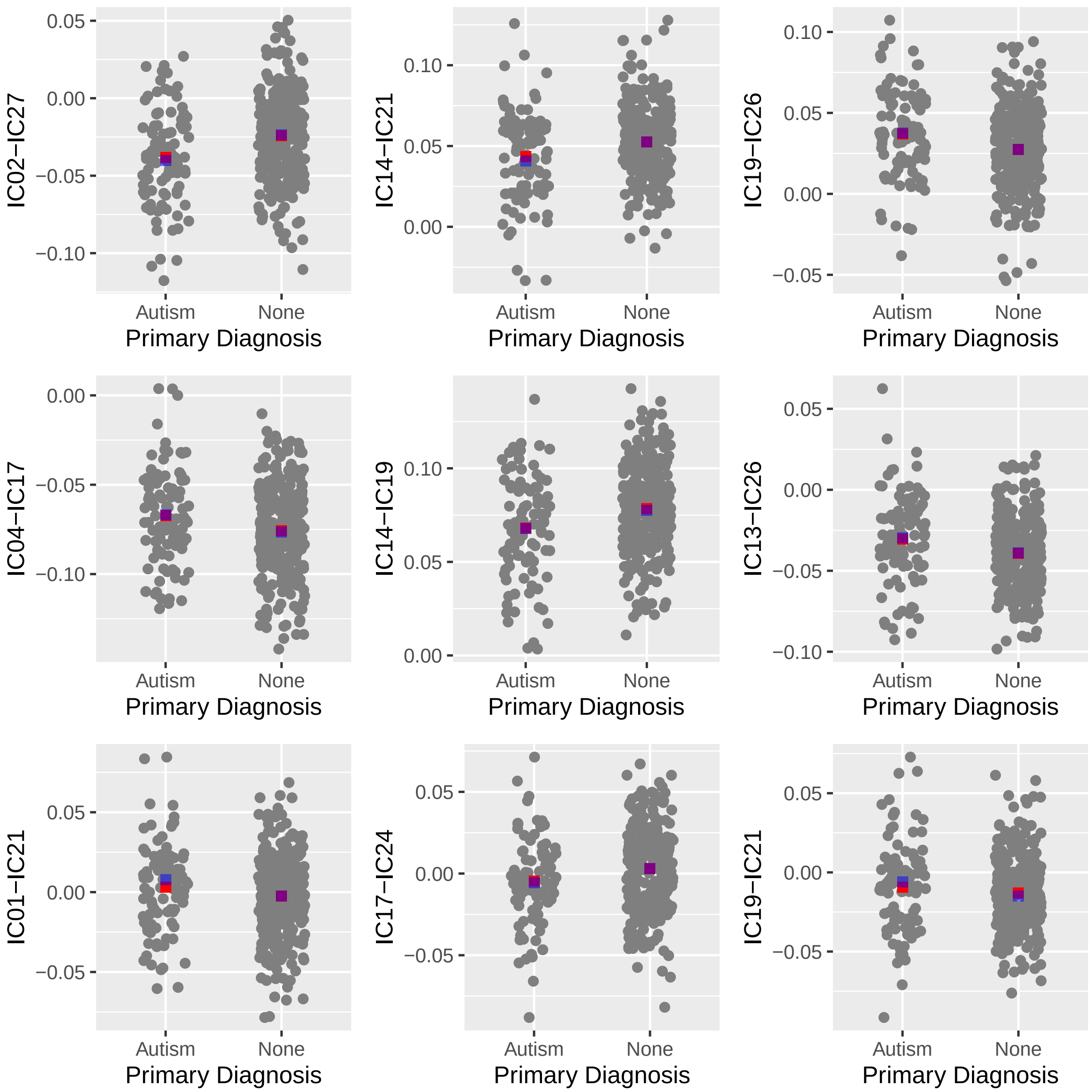


Figure S.4: **Plots of partial correlations, naïve means, and DRTMLE means for each diagnosis group for the nine components with the smallest DRTMLE p-values**. Naïve means appear in red. DRTMLE means appear in blue, with opacity such that means close to each other result in purple. Partial correlations are equal to the adjusted residuals described in the main manuscript Section 2.3.3. These plots indicate that there is large within group variability in the partial correlations. When applied in conjunction with the lenient motion QC, the DRTMLE means are very similar to the naïve means.
